# Embeddings from standardized sorghum leaf images capture variation in disease response that human scoring misses

**DOI:** 10.64898/2026.09.21.753202

**Authors:** Jensina M. Davis, Jonathan Turkus, Sofiya Arora, Karla Montserrat Cuéllar-Perez, Libia Fernanda Gómez-Trejo, Rubén Ruvalcaba-Ramírez, Chidanand Ullagaddi, Xianyan Kuang, Somashekhar Punnuri, Saet-Byul Kim, James C. Schnable

## Abstract

Ordinal scoring of plant disease severity by human raters compresses variation in lesion color, size, and number. Inter-rater variability further complicates comparisons and integrated analyses across environments. We developed a low-cost portable imaging chamber to rapidly image large numbers of leaves under standardized lighting, orientation, and backdrops in the field and employed this system to image more than 11,000 leaves across three states. Embeddings from vision encoders predicted human-assigned disease severity scores. No significant GWAS hits were identified using human-assigned or vegetation-index-based disease severity scores, but GWAS using embeddings identified twelve genomic hotspots controlling leaf appearance. Nine were linked to variation in disease symptom severity. Five hotspots corresponded to previously characterized sorghum genes: all three hotspots not linked to disease and two of the nine that were. Roughly one-third of tested embedding–hotspot associations replicated across at least two states, and twenty replicated across all three. eQTL, PheWAS, and large-effect variant analyses identified single candidate genes with plausible mechanistic links to disease symptom severity for six of the seven hotspots not mapping to characterized genes. These results demonstrate the power of combining scalable, standardized leaf imaging with pretrained image encoders to capture genetically controlled variation in diverse disease symptoms that human ordinal scoring misses.

## 1 Introduction

The global yield of major cereals is reduced by an estimated 17.6–21.0% each year by disease [1]. The scale of these yield losses is forecast to grow as ongoing shifts in growing conditions allow pathogens currently found only in tropical latitudes to extend their range into higher latitudes [2]. Identifying genetic loci conferring resistance to plant pathogens plays an ongoing and critical role in mitigating the impact of disease on crop productivity [3, 4]. Identifying, mapping and confirming disease resistance loci typically requires evaluating hundreds or thousands of plant varieties in studies replicated across multiple environments. Previous GWAS for anthracnose response in sorghum have reported suggestive but often non-overlapping loci, as expected for a trait that exhibits both substantial genotype-by-environment interactions and substantial inter-rater variability [5–11]. In addition, many loci involved in resistance or susceptibility are expected to be segregating at extremely low frequencies, creating challenges for identification and replication, particularly in mapping populations of *<*400 genotypes.

Plant disease responses in field environments are typically quantified via ordinal scores of visual disease severity assigned by human scorers [12–14]. Multiple diseases can be present within the same field study, and human raters can require substantial training to accurately distinguish among diseases as well as between true disease symptoms, leaf damage, and senescence [9]. The requirement for human expert evaluation limits the proportion of field experiments in which disease symptoms are evaluated and reported. In addition, the disease severity scores assigned by different raters to the same plots can vary substantially [9, 10]. The symptoms of foliar (leaf) diseases can vary in lesion size, shape, color, and number, among other factors. Variation in these symptoms is presumably under partially independent genetic control. In some cases specific loci are known to primarily influence a single disease phenotype (e.g., the sorghum *P* locus that controls variation in disease lesion color [15]). Typical ordinal scoring rubrics compress variation in these different phenotypes onto a single axis of disease severity. Differences in the implicit weights raters assign to each component symptom may explain why the same QTL is sometimes detected by one rater’s scores but not others’, or is assigned substantially different effect sizes across raters who evaluated the same plants in the same study [11]. When different raters collect data in different environments, inter-rater variability can be confounded with both environmental effects and genotype by environment interaction (GxE).

Computer vision approaches to quantifying disease severity have been proposed and evaluated as an alternative to address the rater bottleneck in disease phenotyping [16–20]. Images collected from UAV platforms can in some cases be sufficient to predict human-assigned scores of plot-level disease severity [20]. However, these approaches may face challenges in identifying variation in disease-response symptoms among varieties, as image resolution is often insufficient to capture individual lesions or to distinguish specific diseases [21]. Proximal imaging can collect higher resolution images, but, particularly for field-collected images, suffers from challenges in correcting for variation in distance/angle/lighting between images, as well as segmentation of the plant/leaf-of-interest when the background consists largely of other plants [18, 22]. When the challenges of collection variability and segmentation are overcome, computer-vision-based approaches and/or supervised regression models trained to quantify disease severity have the potential to address the challenge of inter-rater variability, as the same indices or trained models can be used to score plant images collected across multiple years and locations. However, supervised models trained on human-assigned disease severity scores replicate the collapse of multi-axis disease phenotypes onto a single axis.

One approach to capture multiple axes of variation in images is to use compressed image representations generated by deep learning models [23, 24]. These representations can be generated in multiple ways including by training autoencoders to describe the variation in a given set of images via a compressed set of latent variables or by passing an image through a pre-trained vision encoder and extracting values from one of its intermediate layers [23, 25]. Autoencoder-based methods are trained directly on the image set of interest, so, at least in principle, latent features should all be informative for variation in that dataset. Embeddings from vision encoders such as DINOv2 [26, 27] and SAM3 [28] benefit from training on internet-scale image datasets, which can yield richer and more general visual representations. However, for any given use case a large proportion of potential embeddings may be non-informative. Previous work has demonstrated the usefulness of latent features extracted from image, hyperspectral, and LiDAR data for studying the genetic control of non-disease phenotypes [23, 24, 29–32].

Here we design and deploy a low-cost imaging chamber that enables proximal leaf imaging under field conditions with consistent lighting, angle, camera distance and clear contrast between leaves and background. We employ this system at research sites in Nebraska, Georgia, and Alabama to collect more than 11,000 images of sorghum leaves in field studies where natural infections of anthracnose (*Colletotrichum sublineola*), a foliar disease that can reduce sorghum yields by *>*70% [33, 34], were observed. We evaluate human scoring, color index, and embedding-based approaches to quantify disease symptoms. Genome-wide association studies conducted using embedding phenotypes for *>*875 sorghum genotypes with 6.4M markers identified hotspots of variation corresponding to both known genes and previously uncharacterized genes belonging to families with known roles in disease resistance. Specific embeddings linked to disease-associated GWAS hotspots reflected a combination of shared signal missed by human-assigned disease severity scores and locus-specific signals which may reflect variation in specific disease symptoms.

## 2 Results

### 2.1 Portable field imaging and vegetation indices quantify leaf health

Eight leaf imaging chambers were fabricated as described in Methods (Fig. 1a). Initial testing determined that training new workers to use the imaging chambers required approximately 10 minutes. After training, individual workers were able to collect between 109 and 196 leaf images per hour. These chambers were employed by separate teams of workers to collect images from three large replicated sorghum field experiments during the 2025 growing season (Fig. 1b) where natural anthracnose (*Colletotrichum sublineola*) outbreaks had been observed. At the time of collection, sorghum plants exhibited a range of disease response and senescence phenotypes (Fig. 1c). Eleven vegetation indices were evaluated for their ability to distinguish healthy and unhealthy leaf tissue across 600 images (Table S1). The Excess Green (ExG) index with a threshold of *−*0.0303 was selected to quantify healthy versus unhealthy leaf tissue (Fig. S1a). The proportion of leaf area classified as unhealthy by this metric had a repeatability of 0.62 across all images collected at the Nebraska field site.

**Fig. 1.**
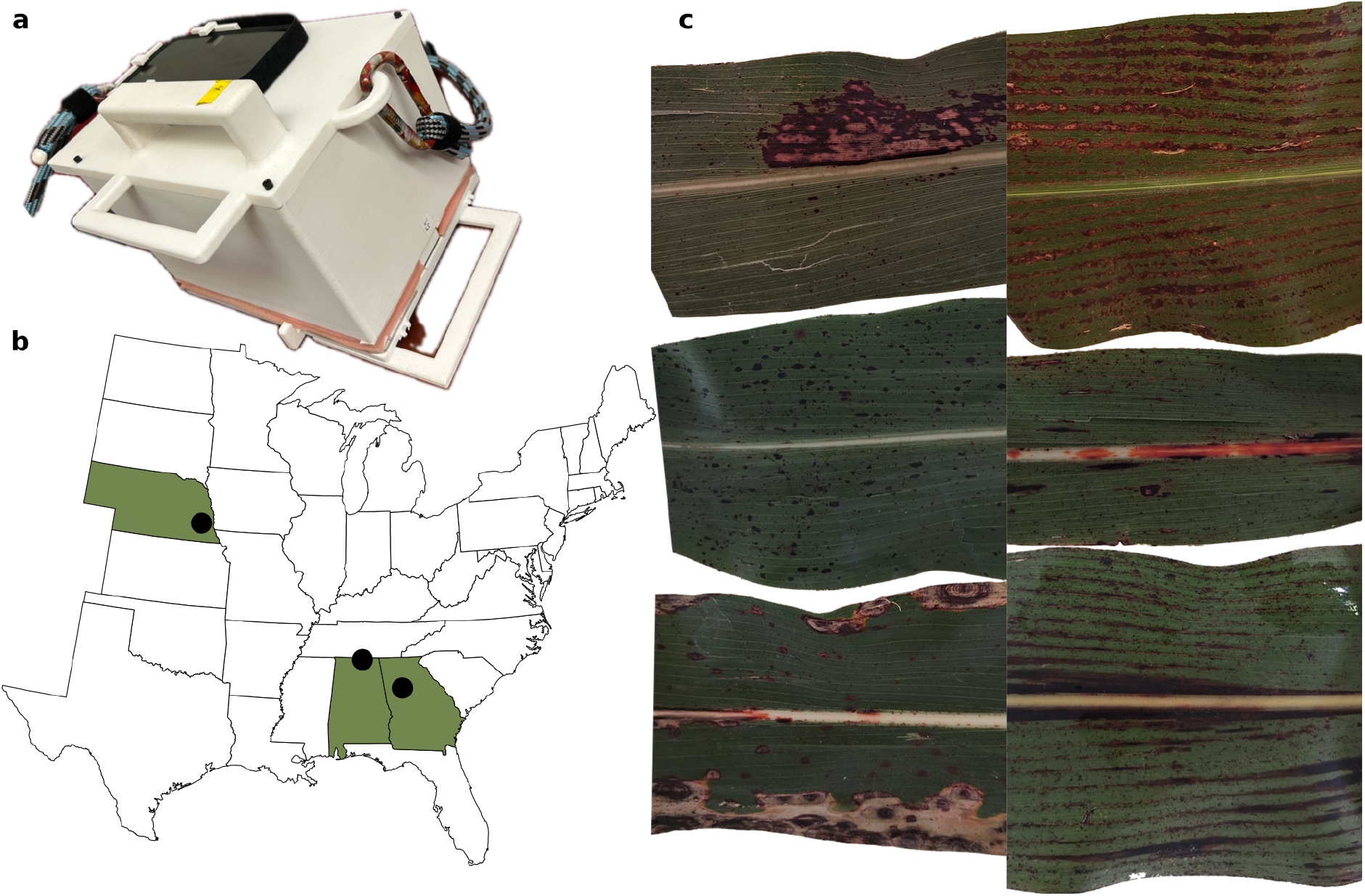
Low-cost, scalable field leaf imaging enables comparison of multiple disease severity indices across environments. **a** Image of portable leaf imaging chamber utilized to collect leaf images. **b** Map of field trial locations where images were collected from sorghum diversity panels. **c** Six example leaves from Nebraska illustrating the diverse appearance of disease responses in sorghum.

On a five-point scale, the mean absolute difference in anthracnose severity assigned by two different human raters (Fig. S1b) was 0.40 at the image level and 0.50 at the plot-mean level. While scores assigned by the two human raters to the same leaf images exhibited substantial disagreement (Fig. S2a), the mean of human-assigned disease severity scores had a repeatability of 0.70. The computationally estimated vegetation-index-based disease severity value, defined as the proportion of all leaf tissue in a given image flagged as unhealthy, was nearly as strongly correlated with the average disease severity score assigned by the two human raters as the scores assigned by the individual raters were with each other (human–VI: Spearman *ρ*^2^ = 0.42 vs. human–human: *ρ*^2^ = 0.45). The relationship between human-assigned disease severity scores and vegetation-index-based disease severity scores broke down in leaves where the vegetation-index-based method classified *>*50% of leaf tissue as diseased (Fig. S2b). Visual examination of these extreme cases determined that many were leaves with substantial amounts of senescent tissue (Fig. S3), and several showed symptoms of a separate, non-anthracnose bacterial disease. Neither the genome-wide association study (GWAS) of human-assigned disease severity scores nor the GWAS of VI-based estimates of the proportion of unhealthy tissue identified signals that exceeded the Bonferroni-corrected multiple-testing threshold.

### 2.2 Embeddings from pre-trained foundation models predict human disease severity scores

We evaluated a range of unsupervised approaches to extract quantitative measurements of image variation from collected leaf images. Latent features from autoencoders trained on the leaf image dataset did not predict variation in human-assigned disease severity scores whether used directly or reduced to 100 principal components (data not shown). Many mean values and/or standard deviations of embeddings extracted from leaf images via either pre-trained DINOv2 or SAM3 vision encoders were correlated with human-assigned disease severity scores. Random forest models trained on these embedding summaries predicted human-assigned disease severity scores (Spearman *ρ*^2^ = 0.60–0.64; Figs. 2a–c, S4). While only minor differences in prediction accuracy were observed between the embeddings generated by the two models and whether mean, standard deviation, or both were employed, including the standard deviation for each embedding improved performance relative to a mean-value-only model for both SAM3 and DINOv2 (Fig. 2a).

**Fig. 2.**
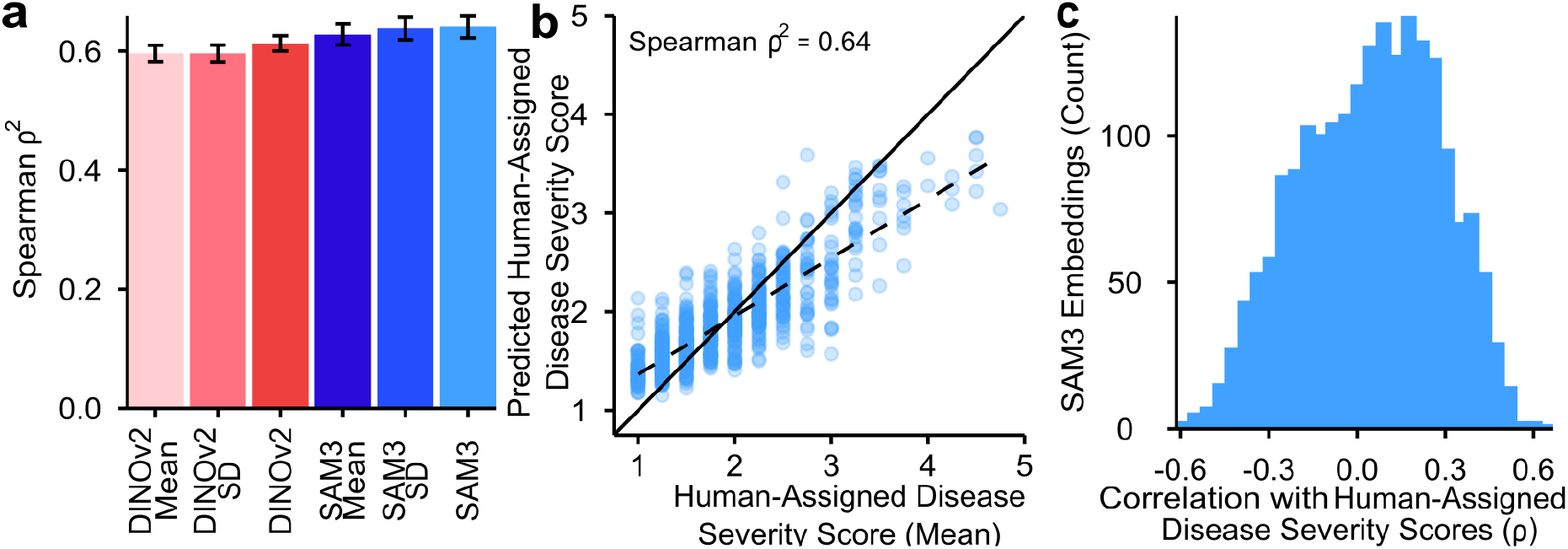
Image embeddings from pretrained models capture disease response in sorghum leaves. **a** Accuracy (Spearman *ρ*^2^) achieved in the Nebraska dataset when predicting human assigned disease scores using different sets of embeddings extracted from either DINOv2 or SAM3. Cross-bars indicate standard error estimated from 5-fold cross-validation. **b** Relationship between the mean disease severity score assigned to images by human raters and the score predicted by a random forest model trained on SAM3 mean+SD embeddings across all five folds of cross-validation. **c** Distribution of observed Spearman correlations between individual SAM3 mean+SD embeddings and human-assigned disease severity scores.

When random forest models were trained to predict the percent leaf area below the ExG threshold from embeddings rather than human-assigned disease severity scores, images with the largest prediction errors tended to contain primarily senescent rather than diseased tissue. This pattern was observed for both SAM3 and DINOv2 and contrasted with low-error images having similar proportions of unhealthy leaf tissue (Fig. S5). SAM3 and DINOv2 embeddings captured varying levels of genetically controlled phenotypes, with repeatabilities in Nebraska ranging from 0.10–0.78 for SAM3 embeddings and 0.00–0.67 for DINOv2 embeddings. SAM3 embeddings that were more strongly correlated with human-assigned disease severity scores were also slightly more repeatable, but this pattern was not observed in DINOv2 embeddings (SAM3: *ρ*^2^ = 0.15, DINOv2: *ρ*^2^ = 5.63 *×* 10*^−^*^5^, Fig. S6).

### 2.3 GWAS identifies embedding hotspots associated with disease resistance and agronomic traits

A total of 747 SAM3 and 610 DINOv2 embeddings were significantly associated with one or more regions of the sorghum genome via GWAS. For both SAM3 and DINOv2, GWAS hits for many different embeddings tended to cluster together in a modest number of hotspots (Fig. 3a). DINOv2 embeddings were somewhat more tightly clustered than SAM3 embeddings: 168, more than one quarter of all embeddings with a GWAS hit, mapped to the 59.9–61.5 Mb region of chromosome 9, which carries one of the three major sorghum height-effect loci, *dwarf1*. We defined 12 leaf-appearance hotspot regions in the sorghum genome, defined as regions consisting of one or more contiguous 100-kb windows containing genetic markers significantly associated with variation in *≥* 10 SAM3 embeddings (Table 1). Applying the same definition using DINOv2 embeddings produced eight hotspots, five of which overlapped with the regions defined via SAM3 embeddings, with the remaining three consisting of clustered but non-contiguous windows on chromosome 6 between 51 and 52 megabases (Table S2). Many image embedding variables mapping to the same hotspots would be consistent with the embedding set containing many redundant variables. However, embeddings associated with the same hotspot typically exhibited only modest correlations with each other (Fig. S7). After controlling for population structure, human-assigned disease severity scores, unhealthy leaf tissue percentage, and the peak marker of a given hotspot, embeddings with GWAS hits in the same hotspot were more correlated with each other than embeddings with GWAS hits at different hotspots (Fig. S8). This is consistent with different embeddings mapping to the same hotspot reflecting, to some extent, variation in specific common traits independent of the specific impact of the locus these embeddings map to and, for that matter, independent of overall disease severity assessed either by humans or by computer vision methodologies.

**Fig. 3.**
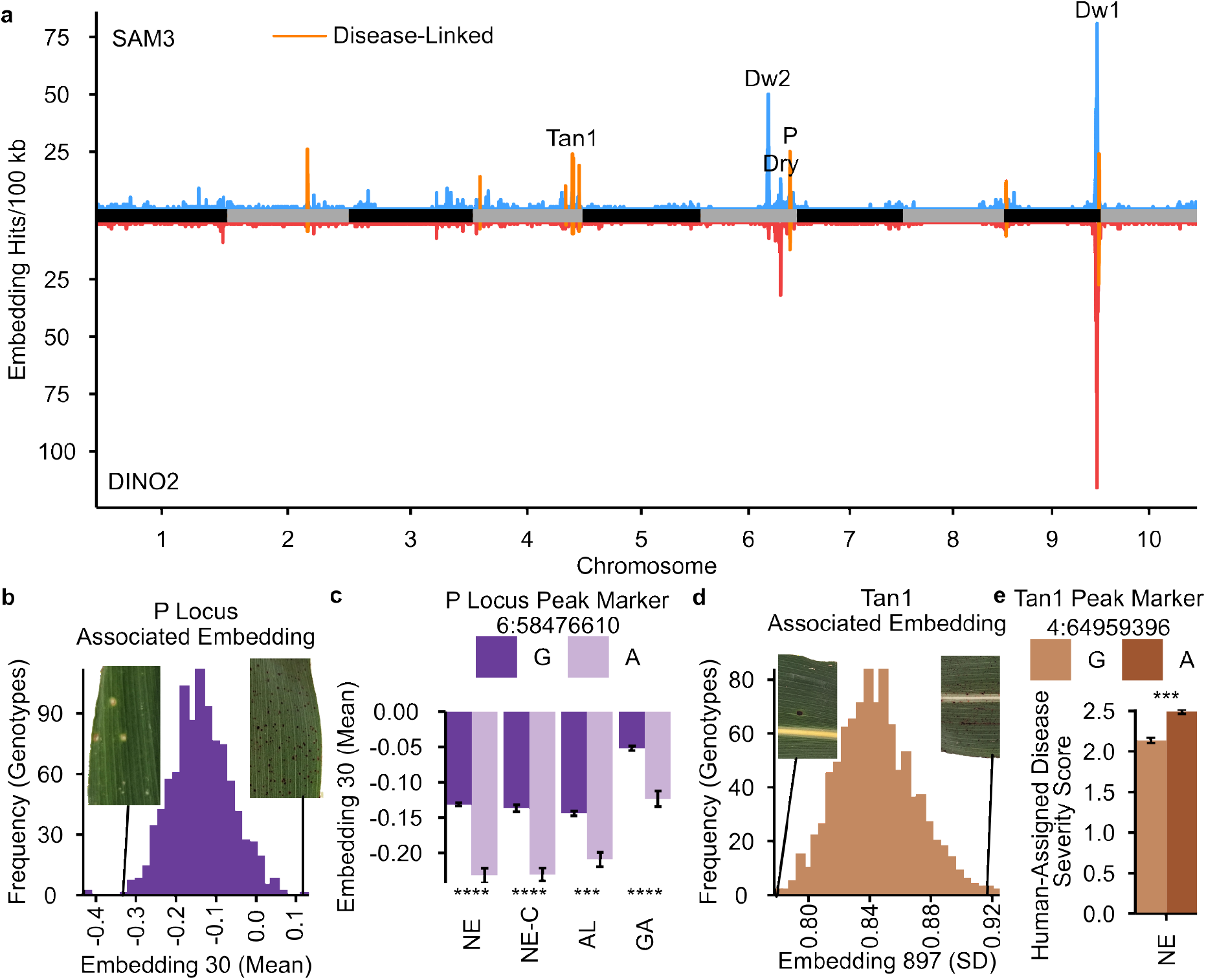
Embedding GWAS hotspots associated with disease response and agronomic traits. **a** Counts of significant SAM3 (top) and DINOv2 (bottom) embeddings in 100-kb windows; orange denotes disease-linked hotspots. **b, d** Nebraska BLUE distributions for SAM3 embedding 30 mean (*P*) and embedding 897 standard deviation (*Tan1*); inset guide lines locate example genotypes. **c** Embedding 30 mean BLUEs by *P* allele. NE, Nebraska; NE-C, Nebraska genotypes shared across all three environments; AL, Alabama; GA, Georgia. Genotype counts (G/A): NE, 849/39; NE-C, 213/15; AL, 256/16; GA, 305/23. **e** Nebraska human-assigned disease severity score BLUEs by *Tan1* allele (*n* = 647 G and 245 A). Bars show means *±* SE. Asterisks denote single-marker likelihood-ratio tests: \**p <* 0.05, \*\**p <* 0.01, \*\*\**p <* 0.001, \*\*\*\**p <* 0.0001.

**Table 1.** SAM3 embedding GWAS hotspots and putative genes. Peak bounds are rounded to 0.1 Mb; marker positions are in base pairs (BTx623 v5). Counts are distinct associated SAM3 embeddings per hotspot. The peak marker has the strongest embedding association within the hotspot. Disease linkage denotes nominal *p <* 0.05 for the peak marker’s association with Nebraska human-assigned disease severity scores.

| Peak | Peak marker | Embeddings | Human score $p$ | Putative Gene |
| --- | --- | --- | --- | --- |
| **Disease-linked** |  |  |  |  |
| Chr02:52.3–52.7 | Chr02:52490664 | 40 | $1.57 \times 10^{-5}$ | *Sobic.002G164900* |
| Chr04:4.7–4.8 | Chr04:4724594 | 14 | $2.90 \times 10^{-5}$ | *Sobic.004G057900* |
| Chr04:60.5–60.6 | Chr04:60556616 | 10 | $4.25 \times 10^{-5}$ | *Sobic.004G230800* |
| Chr04:64.9–65.0 | Chr04:64959396 | 24 | $2.33 \times 10^{-4}$ | *Sobic.004G280800* |
|  |  |  |  | *Tan1* |
| Chr04:65.4–65.5 | Chr04:65447981 | 22 | $1.55 \times 10^{-5}$ | *Sobic.004G286700* |
| Chr04:69.4–69.5 | Chr04:69421678 | 19 | $1.29 \times 10^{-6}$ | Unresolved |
| Chr06:58.2–58.7 | Chr06:58476610 | 30 | $2.39 \times 10^{-4}$ | *Sobic.006G226800* |
|  |  |  |  | *P* |
| Chr09:1.7–1.8 | Chr09:1768703 | 12 | 0.016 | *Sobic.009G019100* |
| Chr09:61.9–62.4 | Chr09:62301540 | 54 | $1.58 \times 10^{-5}$ | *Sobic.009G249900* |
| **Non-disease-linked** |  |  |  |  |
| Chr06:43.5–44.6 | Chr06:43748037 | 75 | 0.418 | *Sobic.006G067700* |
|  |  |  |  | *Dwarf2* |
| Chr06:52.1–52.4 | Chr06:52281164 | 20 | 0.296 | *Sobic.006G147400* |
|  |  |  |  | *Dry* |
| Chr09:59.9–61.5 | Chr09:60857595 | 140 | 0.078 | *Sobic.009G229801* |
|  |  |  |  | *Dwarf1* |

Out of 460 unique combinations of a SAM3 embedding and a GWAS hotspot, 307 were testable and significant (*p <* 0.05) when tested on Nebraska data from only the 230 genotypes shared across all three environments and the genetic marker file used for these analyses. Of these, in 99 cases (32%), the same embedding was significantly associated with the same peak marker within the hotspot in at least one of Alabama and Georgia, and 20 embedding-hotspot associations (6.5%) replicated across both independent environments. Nine of twelve SAM3 embedding GWAS hotspots retained at least three testable and significant embedding-hotspot associations after subsetting to only data from the 230 common sorghum genotypes, and all nine of these hotspots included at least two embeddings whose association with the hotspot replicated in at least one of the two external environments. Five hotspots included at least one individual embedding whose association replicated across all three environments (Table S3).

Five of the twelve SAM3 embedding GWAS hotspots were associated with known, large-effect sorghum genes for which functionally distinct alleles that affect leaf phenotype segregate in our population (Fig. 3a), and nine of the twelve were significantly associated (*p <* 0.05) with variation in human-assigned disease severity scores for the same population of leaf images used to generate the embeddings. The three embedding GWAS hotspots not linked to disease were all cases where a known segregating gene with a large effect on leaf appearance was present in the hotspot. These genes were *Dw2* (*Sobic.006G067700* ; [35]), *Dry* (*Sobic.006G147400* ; [36]), and *Dw1* (*Sobic.009G229801* ; [37]), using sorghum v5 gene model IDs. Among the nine hotspots linked to variation in human-assigned disease severity scores in Nebraska, only four remained significant when evaluated using Nebraska data for only the subset of sorghum genotypes shared across the Nebraska, Alabama, and Georgia field experiments. Of these four, three also exhibited a significant association with human-assigned disease severity scores in at least one of the two independent environments (Fig. S9).

### 2.4 Image embedding hotspots linked to disease are also associated with known genes or plausible candidates

The nine disease-linked leaf embedding GWAS hotspots included two cases that overlapped with well-characterized genes known to be segregating in our population. Variation in a set of 24 SAM3 embeddings was mapped to a tight interval on sorghum chromosome 4 (64.9–65.0 megabases) containing *Sobic.004G280800* (*Tan1*), a WD40 protein first described as a regulator of tannin and anthocyanin biosynthesis in grain [38] and later proposed to regulate 3-deoxyanthocyanidin biosynthesis [39] (Fig. 3d,e). Thirty SAM3 leaf embeddings mapped to a 500-kb interval on sorghum chromosome 6 (58.2–58.7 megabases) containing *Sobic.006G226800*, the causal gene for the sorghum *P* locus [15], which determines whether sorghum leaf lesions develop a purple or tan color (Fig. 3b). The effect of this locus on embeddings was consistent and significant across the complete Nebraska dataset, the overlapping-genotype Nebraska subset, and the Georgia and Alabama field experiments (Fig. 3c). Human experts rated genotypes with the reference (purple) allele at this locus as having significantly higher anthracnose severity than lines with the alternate (tan) allele across all three environments studied (Fig. S9). The *P* locus hotspot exhibited the strongest consistency across environments of any hotspot evaluated, with ten of twenty-five embeddings that were testable and significant in the shared genotype subset of Nebraska data replicating in both the Alabama and Georgia datasets (Table S3).

The remaining seven SAM3 hotspots were not co-localized with a previously characterized gene where alleles with large phenotypic effects were known to be segregating in our population. However, in six of the seven cases, a combination of phenome-wide association studies, additional image analysis metrics, eQTL analysis, and identification of large-effect variants in linkage with the markers identified via GWAS was able to identify candidate genes with plausible or clear potential mechanisms to impact disease resistance and/or leaf appearance. The sole exception was the hotspot located on chromosome 4 between 69.4 and 69.5 Mb, which included GWAS hits for 19 SAM3 embeddings. The lead marker, Chr04:69,421,678:C:A, had a rare A allele (minor-allele frequency = 2.76%; 20 AA and eight CA genotypes among 871 Nebraska genotypes with nonmissing calls). It was not possible to identify a clear candidate gene or potential visible leaf phenotype associated with this locus beyond human-assigned disease severity score. Examination of the individual GWAS results for the 10 SAM3 embeddings that mapped to the Chr4:60.5 Mb hotspot and linkage disequilibrium resolved the hotspot to a smaller 24.8-kilobase interval (Chr4:60,556,616–60,581,417; Fig. S10a). The minor allele of the peak marker, a one-base-pair indel (Chr04:60,556,616:TC:T), was associated with the near abolition of the expression of *Sobic.004G230800* (mean expression = 1.04 TPM for major-allele homozygotes and 0.036 TPM for minor-allele homozygotes; *p* = 1.07 *×* 10*^−^*^6^; Fig. S10b), a gene located 5 kilobases upstream that encodes a UDP-glycosyltransferase whose rice ortholog is associated with changes in stomatal opening [40]. The haplotype associated with the Chr2:52.3 Mb hotspot spanned approximately 310 kilobases (Chr2:52.33–52.64 Mb). The alternate allele for the peak marker (Chr02:52,490,664:GGAGT:G) was associated with an increase in apparent leaf glossiness (*p* = 2.72 *×* 10*^−^*^4^; Fig. S11b) in addition to more severe human-assigned disease severity scores (*p* = 1.57 *×* 10*^−^*^5^; Fig. S11c). The alternative allele was also associated with a reduction in leaf water content in data scored for a subset of the same sorghum population in two different environments (*p* = 0.017 (Michigan 2020), *p* = 1.53 *×* 10*^−^*^4^ (Michigan 2021); Fig. S11g) [41]. The large interval defined for this hotspot based on LD contains nine annotated genes. One of these, *Sobic.002G164900*, located 7 kilobases from the peak marker, encodes a GDSL esterase/lipase whose rice ortholog has been linked to abnormal wax deposition, increased cuticle permeability, and rapid water loss based on analysis of loss-of-function alleles [42]. The Chr4:4.7 Mb hotspot was refined to the interval Chr04:4,712,779–4,746,402 containing six annotated genes. The allele of the peak marker Chr04:4,724,594:G:C associated with greater disease severity was also associated with a substantial reduction in the expression of *Sobic.004G057900* (mean expression = 9.81 TPM for GG and 6.71 TPM for CC; *p* = 7.67 *×* 10*^−^*^7^), a leaf-expressed CYP97B carotenoid hydroxylase inferred to function in lutein biosynthesis based on the functional characterization of orthologous proteins [43] (Fig. S12a–c). Among 29 genes whose expression was screened for association with the peak marker (Chr04:65447981:G:A) of the hotspot, *Sobic.004G286700*, located 5.8 kb distant from the peak marker and encoding a GDSL esterase/lipase (Fig. S11d), had the strongest expression association with the peak marker (*p* = 2.53 *×* 10*^−^*^11^; Fig. S11e). Two linked substitutions within the same codon of *Sobic.004G286700* produce a His-to-Ser substitution at amino acid 277 and are almost entirely linked to the peak marker for the hotspot (*r*^2^ = 0.993). The allele associated with greater human-assigned disease severity scores and higher expression of *Sobic.004G286700* was also associated with a decline in midrib yellowness (*p* = 5.01 *×* 10*^−^*^5^; Fig. S13).

The final two embedding hotspots, both located on chromosome 9, each had substantially stronger evidence supporting specific candidate genes with plausible links to disease response. The Chr9:62.2 Mb hotspot was strongly supported in both the SAM3 (54 embeddings with GWAS hits) and DINOv2 (41 embeddings with GWAS hits) analyses. The hotspot defined by GWAS hits that exceeded a genomewide multiple-testing-corrected threshold of *p ≤* 10*^−^*^7.95^ spanned approximately 500 kilobases, although a tighter 75-kilobase region (Chr09:62,269,891–62,345,242) containing five annotated genes was defined by linkage disequilibrium with the peak marker Chr09:62301540:T:A (Fig. S12d). The expression of two of the five genes in this tighter mapping interval (and none of 19 genes in a more broadly defined region around the mapping interval) was significantly associated with the lead marker: *Sobic.009G249600* (*p* = 2.60 *×* 10*^−^*^5^, higher expression with the high-disease allele) and *Sobic.009G249900* (*p* = 6.24 *×* 10*^−^*^4^, lower expression with the high-disease allele; Fig. S12e,f). *Sobic.009G249600* encodes a DUF179 protein predicted to be chloroplast-localized based on evidence from orthologs in other species. *Sobic.009G249900* encodes a leaf-expressed jasmonate-amino acid ligase orthologous to OsJAR1 [44]. The final embedding GWAS hotspot was also located at Chr09:1.7 Mb. The allele of the peak marker Chr09:1768703:G:T linked to increased disease severity was also associated with a drastic reduction in the expression of *Sobic.009G019100*, a leaf-expressed LysM-domain receptor-like kinase, with median expression declining from 5.4 TPM in the leaves of plants carrying the allele associated with low disease severity to 0.3 TPM in plants carrying the allele associated with high disease severity (*p* = 6.73 *×* 10*^−^*^11^; Fig. S14b–d). We identified a frameshift variant (Chr09:1754173:TTG:T) within the LysM-domain receptor-like kinase that was not in linkage with the peak marker. However, this potential second allele exhibited a significant association with ExG-estimated leaf damage (*p* = 0.0125), which modestly increased in significance after conditioning on the peak marker; however, it must be noted that the potential second allele was not significantly associated with human-assigned disease severity scores. The allele associated with reduced disease severity and increased LysM-domain receptor-like kinase expression was significantly associated with a reduction in yield-related phenotypes scored across three of six environments tested (Michigan 2020 panicle dry mass *p* = 1.61 *×* 10*^−^*^4^; Michigan 2021 panicle dry mass *p* = 0.00202; Nebraska 2020 grain mass per panicle *p* = 0.0388; Fig. S15) and directionally consistent but individually non-significant effects in two additional environments (Nebraska 2021 and Nebraska 2023). A modest positive impact on grain mass per plant was observed only in Nebraska 2025, in the same anthracnose-infected field trial in which disease severity was assessed.

## 3 Discussion

Meeting the need for an improved understanding of plant disease responses, and for the discovery and characterization of new disease resistance loci necessitated by the ongoing emergence and spread of plant pathogens, requires more and better data on plant disease symptoms collected from larger and more numerous field studies. As discussed above, human scoring of disease severity presents a number of logistical and consistency challenges. However, even when expert human raters conduct visual assessments of disease severity for large numbers of replicated plants from large diversity panels, identifying disease resistance loci is not assured. In our study, two raters with graduate training in plant pathology scored 6,026 images (after quality control exclusions) representing 891 genotypes with high-density markers from whole genome resequencing for anthracnose disease severity, yet no genetic associations passed multiple testing correction (Fig. S16a,b).

While computer vision approaches to scoring disease based on vegetation indices can address the challenges of limited availability of skilled human labor and inter-rater variability, they can struggle to distinguish between different diseases, or between disease and dead tissue resulting from mechanical damage or senescence. In our study, thresholding on the ExG vegetation index reliably separated healthy leaf tissue from unhealthy tissue (Fig. S1) but was unable to differentiate between senescence and disease symptoms (Fig. S3). GWAS conducted using untransformed estimates of disease severity from ExG produced substantially inflated genome-wide results driven by extreme values from largely senesced leaves (Fig. S16c,d), and after a logit transformation to address these extreme values, no genetic associations passed multiple testing correction (Fig. S16e,f). In contrast, image embeddings generated using pretrained vision encoders identified multiple hotspots of leaf appearance variation, including known genes altering either overall leaf appearance (e.g., *Dwarf1*, *Dwarf2*, and *Dry*) or disease response and symptoms (e.g., *P* and *Tan1*) (Fig. 3, Table 1).

Both the *P* and *Tan1* GWAS hotspots were significantly associated with variation in human-assigned disease severity scores. Both loci play roles in determining the identity and quantity of flavonoid molecules, including the 3-deoxyanthocyanidin phytoalexins. These phytoalexins are visible pigments with potential defensive functions against plant pathogens [15, 39, 45, 46]. However, in contrast to the expectation that more pigment would provide more resistance to disease, the allele of the *P* hotspot associated with a functional copy of the flavanone 4-reductase encoded by *P* and the accumulation of 3-deoxyanthocyanidins [15] was also linked to significantly higher human-assigned disease severity scores (Figs. 3, S9). This pattern is instead consistent with human scorers interpreting dark purple lesions as indicating more severe disease than pale tan lesions. Future work could test whether this is the case using generated sorghum leaf images with the same number and size of lesions but different lesion colors.

Embeddings associated with disease-linked hotspots shared information not explained by association with the genomic regions themselves or by human-assigned and computationally estimated metrics of disease severity. This was particularly true of embeddings associated with the same hotspot, although it was also observed for embeddings associated with different hotspots (Fig. S8), indicating that the image embeddings captured shared phenotypes of disease response that are not captured by human-assigned disease severity scores, both broadly and specific to certain loci. Embeddings associated with loci not linked to human-assigned disease severity scores also retained limited shared information both within and across hotspots (Fig. S8), indicating that the image embeddings capture a shared latent phenotype. The interpretation of GWAS results from leaf embeddings requires substantially more investigation than the interpretation of GWAS results from human-assigned disease severity scores. Three-quarters of the embedding GWAS hotspots identified in this study exhibited an association with human-assigned disease severity scores, but in the absence of human-assigned disease severity score data for the same field study, separating disease-linked signals from other, non-disease-related leaf appearance signals would be more challenging. ExG or other index-based metrics have the potential to partially substitute for human scores in identifying which loci are associated with disease, but with the same caveat described above that such metrics may confound disease, senescence, and mechanical damage. Individual marker-trait associations or phenome-wide association studies conducted using additional color or appearance features extracted from the same leaf images used to generate embeddings, and/or published trait data collected from the same population [47], also showed promise as approaches for understanding the phenotype or mode of action underlying specific leaf embedding GWAS hotspots. Perhaps the most striking example of this was provided by the Chr09:1.7 Mb hotspot (LysM-RLK), where the comparatively rare allele (39 of 891 Nebraska 2025 genotypes) associated with increased disease severity in Nebraska 2025 was linked to a significant increase in grain yield in multiple other environments, but was yield-neutral in our field under significant anthracnose disease pressure (Fig. S15), consistent with a growth/defense tradeoff.

Ultimately, this work was only possible because we were able to collect large numbers of images of sorghum leaves across multiple environments using a technology that ensured leaf images were easily segmented and comparable in resolution, lighting, and orientation. The portable imaging chamber described in this study requires less than ten minutes of training to use, and after training, individual workers collected between 109 and 196 leaf images per hour. The chamber was employed by researchers in Alabama and Georgia without any face-to-face interaction with the original design team. The original bill of materials to construct the chamber totaled roughly $250 (Information S1). However, approximately 60% of this cost was the color calibration card; substituting a low-cost alternative reduces the chamber cost to approximately $100.

This study sought to address two barriers to plant pathology research: the necessity of relying on scarce, expensive, and variable human raters, and the use of a single axis to assess disease severity when multiple facets of disease may be under partially independent genetic control. The use of a low-cost imaging chamber to collect many thousands of directly comparable images had the potential to address both challenges. A single rater is able to assess and score disease for thousands of images across multiple environments in a semi-random order (Information S3). While this approach was still unable to identify significant genetic signals, we were able to identify twelve leaf appearance hotspots using embeddings extracted from the same images by pretrained image encoders. Many embedding-hotspot associations replicated across environments (Table S3), and because the imaging setup and encoder can be held constant, the same embedding phenotypes can continue to be quantified in additional field studies in future years. Almost half of these hotspots corresponded to known segregating genes with impacts on leaf appearance. Among the other seven, eQTL and large-effect variant analyses allowed us to identify genes with plausible, specific mechanisms that might alter disease susceptibility and response, in a number of cases supported by associations with additional traits quantified from the same leaf images or by PheWAS conducted across a large panel of traits scored in the same population in multiple environments. These results illustrate the potential of pairing rapid, standardized, and low-effort protocols for in-field imaging with advanced vision encoders to better understand and dissect the genetic basis of plant responses to disease.

## 4 Methods

### 4.1 Field experiments

Three sorghum field experiments were conducted as part of this study. The Lincoln, Nebraska experiment consisted of 968 genotypes drawn from 1) the Sorghum Association Panel (SAP) [48], 2) the Sorghum Diversity Panel (SDP) [49], and 3) varieties with expired Plant Variety Protection, grown in a randomized complete block design with two total blocks and a repeated check genotype at the University of Nebraska-Lincoln’s Havelock Research Farm (40.86*^◦^*N, 96.60*^◦^*W). A total of 2,100 one-row, 1.52-meter experimental plots with 0.76-meter row spacing were planted on June 10, 2025. In Fort Valley, Georgia, 692 genotypes drawn from the SAP and the Sorghum Bioenergy Association Panel (BAP) [50] were planted on June 13, 2025, with each association panel planted in a randomized complete block design with two blocks, for a total of 1,384 one-row, 4.27-meter plots with 0.91-meter row spacing at Fort Valley State University’s New Farm (32.56*^◦^*N, 84.31*^◦^*W). In Hazel Green, Alabama, 338 genotypes comprising a subset of the SAP were grown in a randomized complete block design with two blocks per nitrogen treatment level (112 and 202 kg N/ha) at Alabama A&M University’s Winfred Thomas Agricultural Research Station (34.90*^◦^*N, 86.56*^◦^*W). Each block comprised 432 two-row, 1.83-meter plots with 0.88-meter row spacing (the 338 genotypes plus check and fill plots), for a total of 1,728 plots planted on April 9, 2025.

Phenotypic data from 6 field trials in Alabama, Michigan, and Nebraska from 2020 – 2023 containing subsets of genotypes grown as part of this study and additional traits collected from the Nebraska 2025 trial, from which images described in this study were collected, were aggregated for a total of 121 unique trait x environment combinations across 48 unique traits (Table S4) and utilized for PheWAS.

### 4.2 Portable imaging chamber design and construction

We designed a portable imaging chamber to standardize the camera distance and angle relative to individual leaves in a field environment and to provide relatively consistent lighting conditions. The imaging chamber exterior is constructed from four components 3D-printed using Bambu Lab X1-Carbon printers with white PLA filament to maximize light diffusion and to provide a high-contrast background against the leaf surface. STL design files and a comprehensive parts list are provided in Supplemental Information S1. Briefly, the handle is glued to the phone mount, which is attached to the chamber sides with heat-set inserts and M3 screws, washers, and lock washers (hereafter M3 hardware). The chamber sides are joined to the imaging canvas by a hinge secured with heat-set inserts and M3 hardware. The imaging canvas contains an inset in which a Calibrite ColorChecker Classic Nano card is inserted beneath a plexiglass sheet that is secured with M3 hardware. A handle is affixed to the underside of the imaging canvas using the same M3 screws that secure the plexiglass, reinforced by heat welding with wire mesh. A pair of magnets secured with super glue—one on the bottom center of the chamber-sides perimeter and one on the upper center of the imaging-canvas perimeter—ensures closure opposite the hinge. Polyurethane foam is attached with double-sided tape to the remaining perimeter area of the imaging canvas and the chamber sides to minimize light penetration through their junction point. Artificial lighting is provided by a battery-powered LED light strip (BestLuz BSL-003C) placed around the bottom perimeter of the interior chamber sides. A Galaxy A16 smartphone is held on the phone mount by a hook-and-loop fastener strip attached with adhesive backing and M3 hardware. Carabiners attach a neck strap to side hooks on the phone mount.

### 4.3 Image collection

A detailed written protocol for image collection, including a video demonstration, is provided in Supplemental Information S2. Briefly, a leaf with little or no mechanical damage was placed in the imaging chamber with the adaxial surface facing up, oriented so that the leaf extended beyond the sides of the chamber and did not obscure the color calibration card. The flag leaf and lower canopy leaves were excluded when selecting leaves for imaging. A target of three images of separate leaves was collected per plot. In some cases fewer leaves were imaged when fewer than three plants with intact leaves were present. In a small number of cases, more than three leaf images were collected. A 3060 *×* 4080 RGB image was captured and associated with the plot using a photo trait in Field Book [51]. In Nebraska, 6,147 images were collected on September 8 and 9, 2025, from 1,985 plots. In Georgia, 3,695 images were collected October 1–10, 2025, from 1,231 plots. In Alabama, 1,735 images were collected October 15–24, 2025, from 578 plots.

### 4.4 Image segmentation and pre-processing

Binary masks for each individual image were generated using a computer vision pipeline implemented in OpenCV [52]. Two flood fills based on seeds above and below the typical position of leaves were used to identify white background pixels. After the subtraction of these pixels, the largest connected component which extended to both the left and right edges of the image was evaluated, rejecting cases where this component consisted of fewer than 750,000 pixels (~6% of image area) or more than 7.5 million pixels (~60% of image area) and where this component extended to the region containing the color correction card. Images where the largest connected component failed any of these criteria were excluded from downstream analysis. The final leaf mask consisted of this largest connected component after the removal of the region within 300 pixels of the left image border and 100 pixels of the right image border as these regions sometimes included color variation introduced by exterior light leaking into the imaging chamber. The first two principal axes of each leaf mask were estimated from the x and y coordinates of pixels within the mask. Overlapping crops of 2016 *×* 2016 pixels were generated parallel to the leaf axis. The first crop was generated at the minimum position along the principal axis where all four corners fell within the raw image and the process continued in step sizes of 500 pixels until a candidate crop would extend beyond the raw image. This process typically generated two crops per image.

### 4.5 Human scoring of anthracnose severity

Human expert scoring was conducted on an ordinal scale of disease severity from 1–5 in 0.5 step increments (Fig. S17) using a custom web app that displayed individual images (Information S3). Scoring was conducted by two human raters with graduate training in plant pathology. When two raters scored the same image, the average of both raters’ scores was employed for downstream analyses. All Nebraska images passing segmentation-based checks were scored by humans, as was a subset of images selected based on ExG quintiles in the Alabama and Georgia environments.

### 4.6 Vegetation index thresholding

Non-leaf pixel values were set to zero using the binary masks described above. A random subset of 600 images from all three field sites was initially used to evaluate candidate vegetation indices and identify a pixel-level threshold (Table S1; [53, 54]). Candidate indices were selected based on visual comparison of disease overlays generated from segmented leaf images. Among the vegetation indices evaluated, NGRDI and ExG performed well and exhibited roughly equivalent performance in separating healthy and diseased tissue during initial testing. Further threshold selection for pixel-level classification of unhealthy tissue for these two indices was based on three criteria: (i) distributions of pixel-wise vegetation index values visualized as density histograms; (ii) visual agreement between disease overlays and observed disease symptoms; and (iii) manual evaluation of unhealthy-tissue overlays against the corresponding raw images. The ExG index [55] at a threshold of -0.0303 was selected for the downstream analysis based on consistent visual separation between unhealthy/diseased and healthy tissue. ExG scores employed in downstream analyses are the logit-transformed percentage of leaf pixels falling below the selected ExG threshold within a given leaf mask.

The glossiness of each leaf was defined as the fraction of leaf pixels with a brightness exceeding the leaf’s own mean plus two standard deviations. Values were averaged within genotypes, and two genotypes with extremely low gloss values (*<*0.025) were excluded from downstream analyses.

### 4.7 Generating embedding scores

Embedding values were generated using two pre-trained models: DINOv2 (dinov2 vitl14 reg) [26, 27] and SAM3 (facebook/sam3, applied via the Hugging Face transformers Sam3Model and Sam3Processor) [28]. Individual image crops were resized from 2016 *×* 2016 to 1008 *×* 1008 via cv2.resize(…, interpolation=cv2.INTER AREA). For DINOv2 each channel was normalized to have a mean and standard deviation of 0.5 while normalization within SAM3 was conducted via Sam3Processor. SAM3 patch tokens were extracted from the vision encoder’s last hidden state and DINOv2 patch tokens were extracted from x norm patchtokens. The mean and standard deviation of each of the 1,024 patch-token dimensions across all patches were calculated for each model, resulting in a final set of 2,048 embeddings per model.

### 4.8 Prediction of disease scores via random forest

Random forest models were trained on per-image embedding values generated by averaging values per crop when multiple crops were present per image. Images were split into five folds for cross-validation in a genotype-aware fashion (i.e., all images collected from the same sorghum genotype were assigned to a single fold to minimize potential data leakage). Embeddings were standardized (z-scaled) within each cross-validation fold using statistics estimated on the training partition and applied to the heldout partition. Within each fold, hyperparameters were optimized via random search on the training set using 100 iterations and three-fold internal cross-validation. Feature importances and test set predictions were saved for each test fold. Feature importances were calculated as the mean decrease in squarederror impurity reported by the random forest fit on each training fold and averaged across the five cross-validation folds.

### 4.9 Genome wide association studies (GWAS)

GWAS was conducted using a set of markers generated via alignment of whole genome resequencing of 925 sorghum lines including 891 present in the Nebraska experiment and 274 present in the Alabama experiment and 330 present in the Georgia, respectively, to the BTx623 v5 reference genome [56]. The resequencing data used to generate this marker set included a previously published set of Illumina sequencing data for the SAP panel [57] and additional whole genome resequencing conducted via Ultima for the remaining lines. This marker set comprised 6,422,975 markers.

Trait values were averaged to plot-level means and winsorized at the 1st and 99th percentiles prior to calculating best linear unbiased estimators (BLUEs) via lme4 [58]. Within a single environment, each trait was modeled as

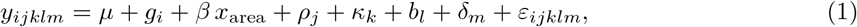

where *µ* is the intercept, *g_i_* is the effect of genotype *i* (fit as a fixed effect), *x*_area_ is the *z*-scored leaf mask area with coefficient *β*, and *ρ_j_*, *κ_k_*, *b_l_*, and *δ_m_* are random effects of field row, column, block, and imaging device, respectively, each distributed *N* (0*, σ*^2^). Estimates of variance attributable to each experimental design factor within a single environment were generated via the same model with the modification that genotype was fit as a random rather than a fixed effect. Repeatability values within environments are *H* = *v_g_/*(*v_g_* + *v_e_/r̅*) where *v_g_* denotes the genotypic variance component, *v_e_* denotes residual variance, and *r̅* is the *harmonic mean* of the number of replicated observations per genotype. For the estimated proportion of diseased leaf pixels from ExG calculations, image-level values were logit-transformed before calculating plot-level means and BLUEs; proportions were clipped to [5 *×* 10*^−^*^5^, 1 *−* 5 *×* 10*^−^*^5^] before transformation. An otherwise identical GWAS using untransformed ExG values was conducted for comparison (Fig. S16). GWAS was conducted using the mixed linear model likelihood ratio test method implemented in PANICLE (https://github.com/jschnable/PANICLE) using leave-one-chromosome-out kinship matrices and 5 principal components calculated from the genetic marker data and Z-score transformed leaf mask area and flowering time as additional covariates. Associations were considered statistically significant if they exceeded a threshold of *−* log_10_ *p ≥* 7.95 corresponding to a p-value of 0.05 adjusted by an estimated 4,446,367 effective independent tests.

Hotspots of associations with embeddings were identified using fixed, non-overlapping 100-kb bins. Hotspots separated by less than 100 kb were merged. The peak marker for a given hotspot region was defined as the marker most significantly associated with any single embedding within that hotspot.

### 4.10 Hotspot characterization and candidate gene investigation

Expression data employed for eQTL analyses were sourced from mature leaf tissue measured in a 2021 field experiment [59]. The association between expression of a given gene, expressed in units of transcripts per million (TPM), and genotype at the marker of interest was evaluated using a single-marker likelihood ratio test in PANICLE with leave-one-chromosome-out kinship and with five genetic-marker PCs, flowering time, and leaf area included as covariates. Other tests of association between specific markers and specific traits were conducted using the same single-marker test approach in PANICLE.

To estimate partial correlations between continuous variables, a least squares model with rank-transformed values for all model terms was fit separately for each variable in the pair including the first 5 principal components from the genetic marker data and human-assigned disease severity score and logit(Percent Unhealthy Leaf Tissue) BLUE values. For partial correlations of embeddings associated with the same hotspot, the alternate allele dosage at the lead marker was additionally included as a covariate. Spearman correlation of the residuals from the models for each variable was estimated. Partial correlations were estimated for all pairs of embeddings associated with the given hotspot(s) extracted from both SAM3 and DINOv2. Significance of differences between partial correlations was tested with a Wilcoxon rank-sum test with continuity correction.

## Acknowledgements

The authors thank Waqar Ali, Ozgur Altundas, Sophie Rohlfing, Sofiya Arora, Harshita Mangal, Zhongjie Ji, Hadiya Kounsar, Sawyer Johnson, Manoj Kumar Sangireddy, Surakshya Ghimire, Janak Adhikari, Asha Colvard, Xinhua Xiao, Blake Long, Abiodun Adeniyi, Friday Zakari, McKinley Dunford, Camille McGowan, Ramel Woodard, and Laron Foster for their assistance in collecting leaf images.

## Author contributions

JMD, JT, and JCS conceived of the project. JMD, SBK, and JCS designed experiments. JMD, JT, SA, KMCP, LFGT, RRR, CU, XK, and SP conducted experiments and generated data. JMD, SA, KMCP, and JCS analyzed the data and visualized and interpreted the results. JMD and JCS drafted the paper. All authors reviewed, edited, and approved the final manuscript.

## Competing Interests

JMD, JT, and JCS are inventors on U.S. Patent Application 19/791,214, “Portable leaf imaging chamber and image analysis methods for field phenotyping.” JCS has equity interests in Data2Bio and Dryland Genetics and has performed paid work for Alphabet. The authors declare no other competing interests.

## Generative AI use statement

The authors used AI models, including ChatGPT-5.5, ChatGPT-6.0, Claude Opus 5, Claude Sonnet 5, and Claude Fable 5.0/5.1, to assist in drafting code for data analysis and visualization and to identify spelling and grammatical errors. The authors reviewed and evaluated in detail all results produced using AI-assisted code and take full responsibility for the accuracy of the work presented in this paper.

## Funding

This work was supported by the Nebraska Corn Board Presidential Chair Endowment fund at the University of Nebraska-Lincoln and the National Science Foundation under grant IOS-2412930. Field experiments at Fort Valley State University were supported by the USDA-NIFA under grant 336172. Data collection and student training at Alabama A&M University were supported by the U.S. Department of Energy, Office of Science, Office of Biological and Environmental Research, under Grant No. DE-SC0024615. JMD is supported by the National Science Foundation Graduate Research Fellowship Program, Grant No. 2034837. This work was completed utilizing the Holland Computing Center of the University of Nebraska, which receives support from the UNL Office of Research and Innovation, and the Nebraska Research Initiative.

## Data Availability

Raw image and genomic marker data is submitted and in queue for curation and release at the time of submission and will be available at https://doi.org/10.5061/dryad.cfxpnvxpb. All analysis and data visualization code utilized in this manuscript is available at https://github.com/jschnable/SorghumLeafEmbeddings/. STL files for the portable leaf imaging chamber are available in Information S1.

## 5 Supplementary information

**Fig. S1.**
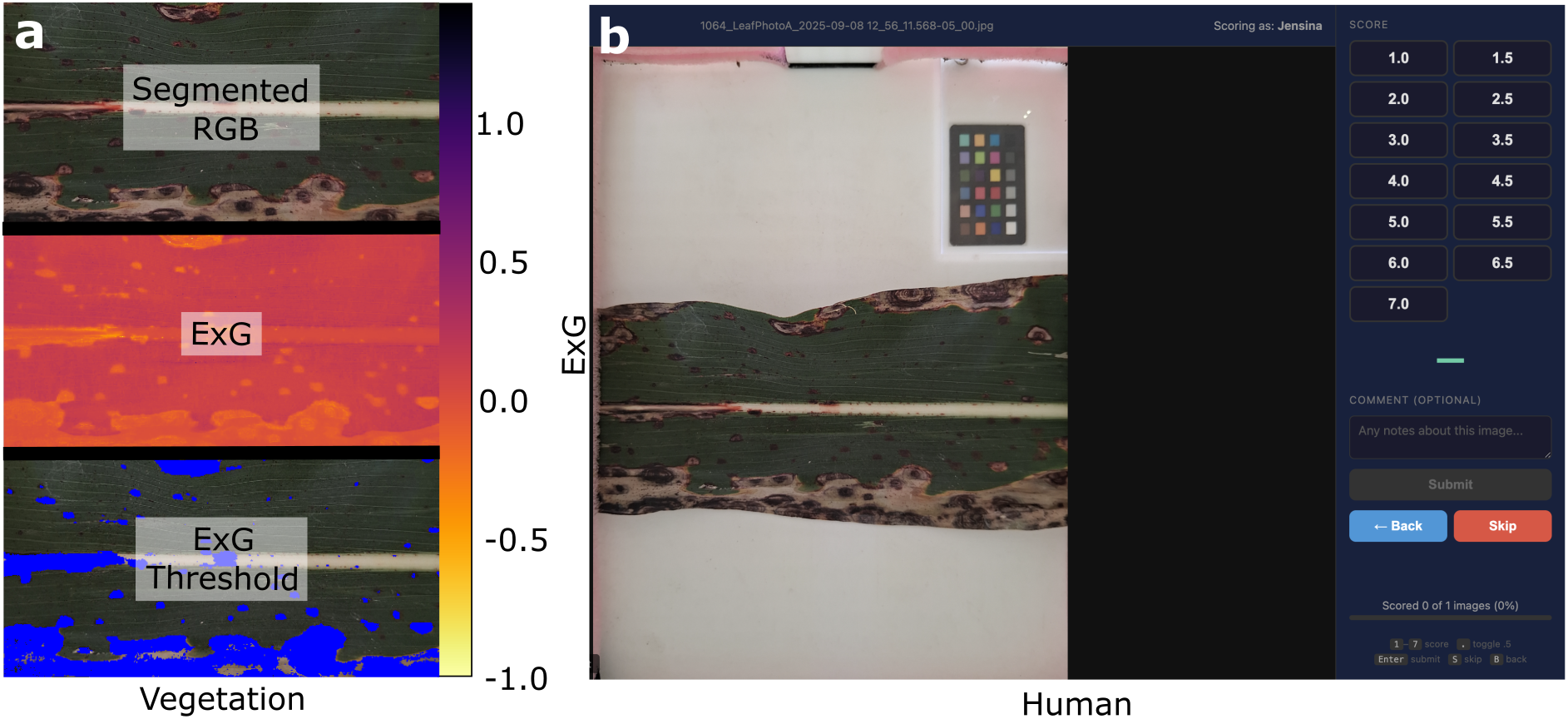
Disease-scoring workflows. **a** ExG thresholding of segmented leaf images. **b** Web interface for human-assigned disease severity scoring.

**Fig. S2.**
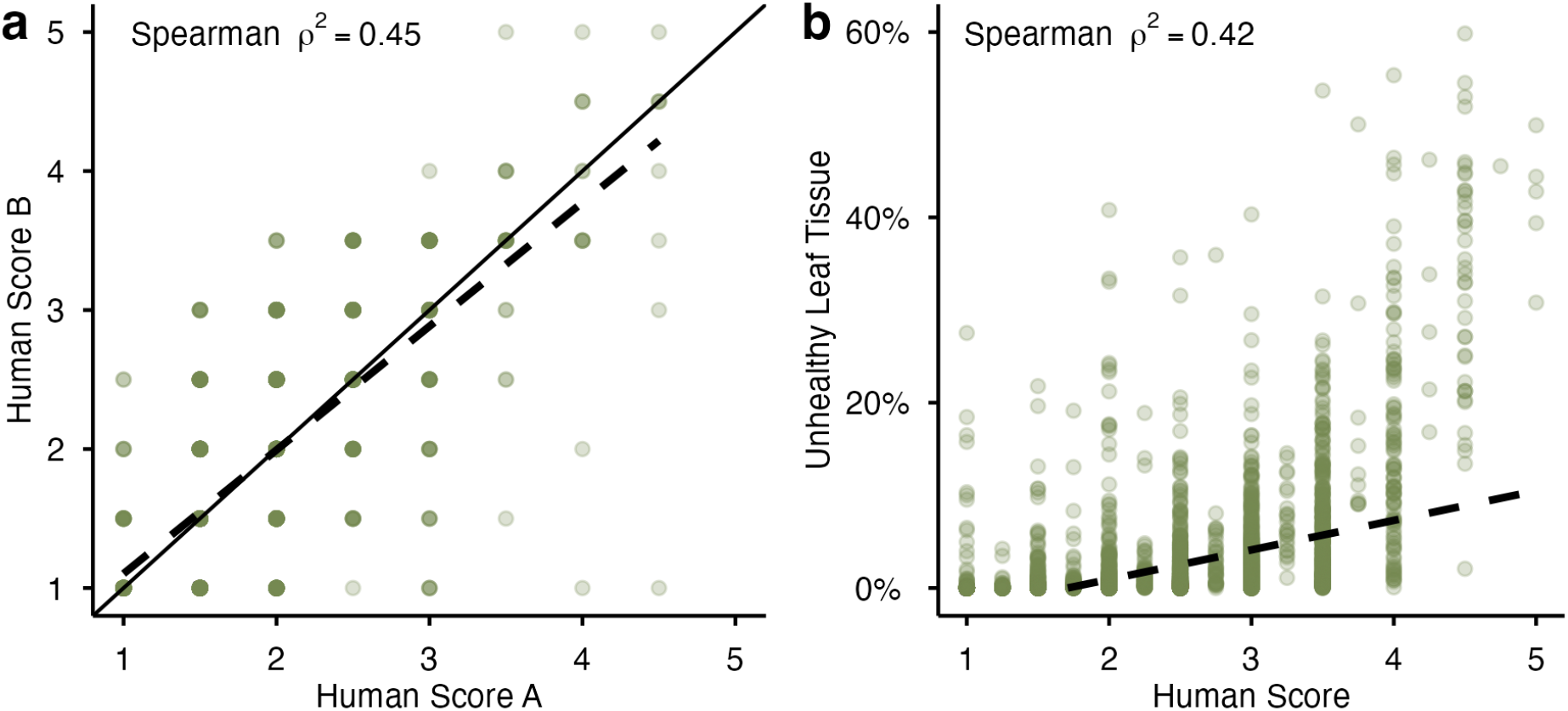
Agreement between disease-scoring approaches. **a** Independent human-assigned disease severity scores of 999 images. **b** Mean available human-assigned disease severity score versus ExG-estimated unhealthy leaf area. Dashed lines show linear fits; the solid line in **a** indicates equality.

**Fig. S3.**
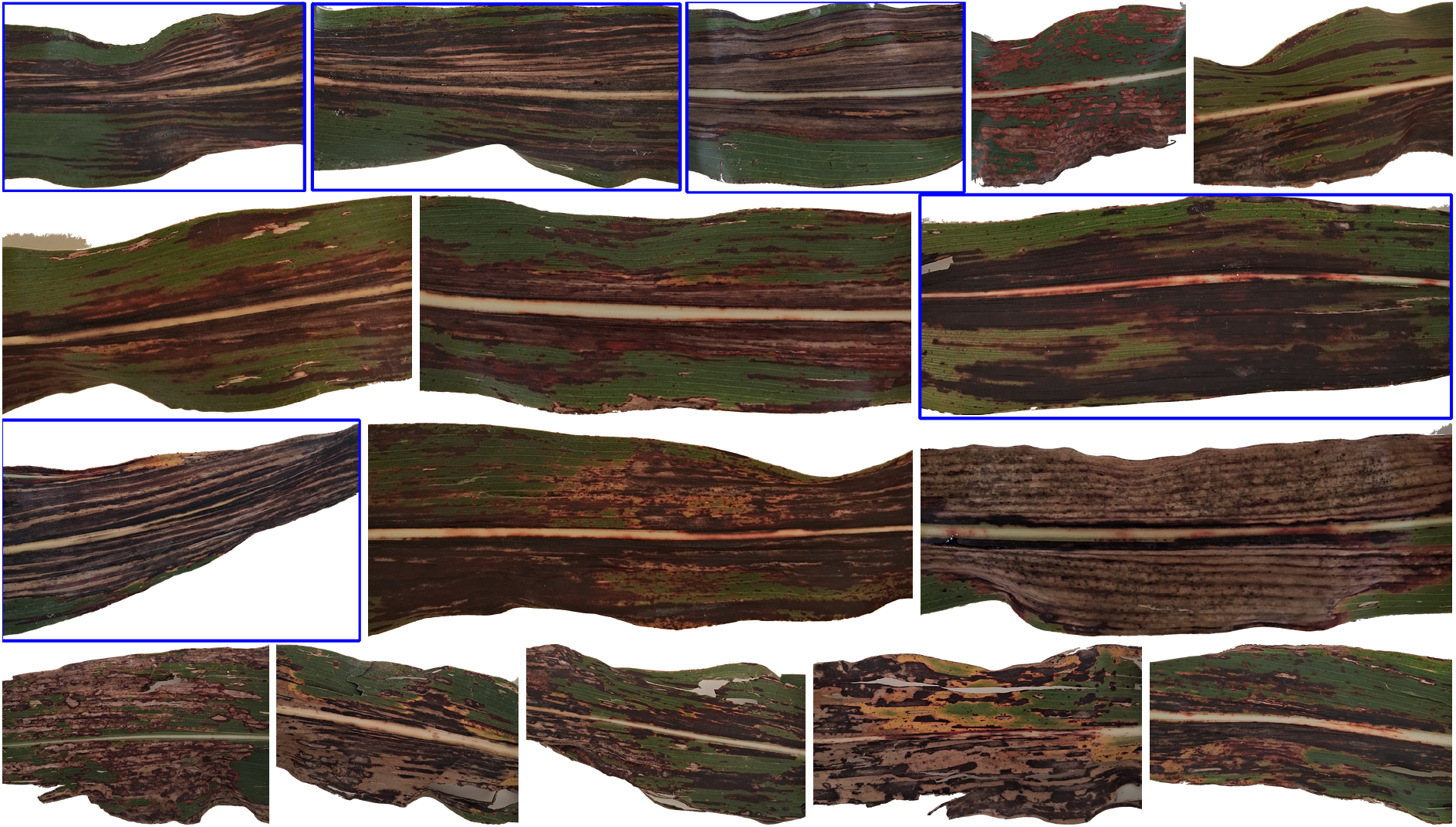
Nebraska leaf images with *>* 50% of leaf area below the ExG threshold. Blue rectangles indicate leaves exhibiting symptoms of infection by a bacterial pathogen rather than by *Colletotrichum sublineola*.

**Fig. S4.**
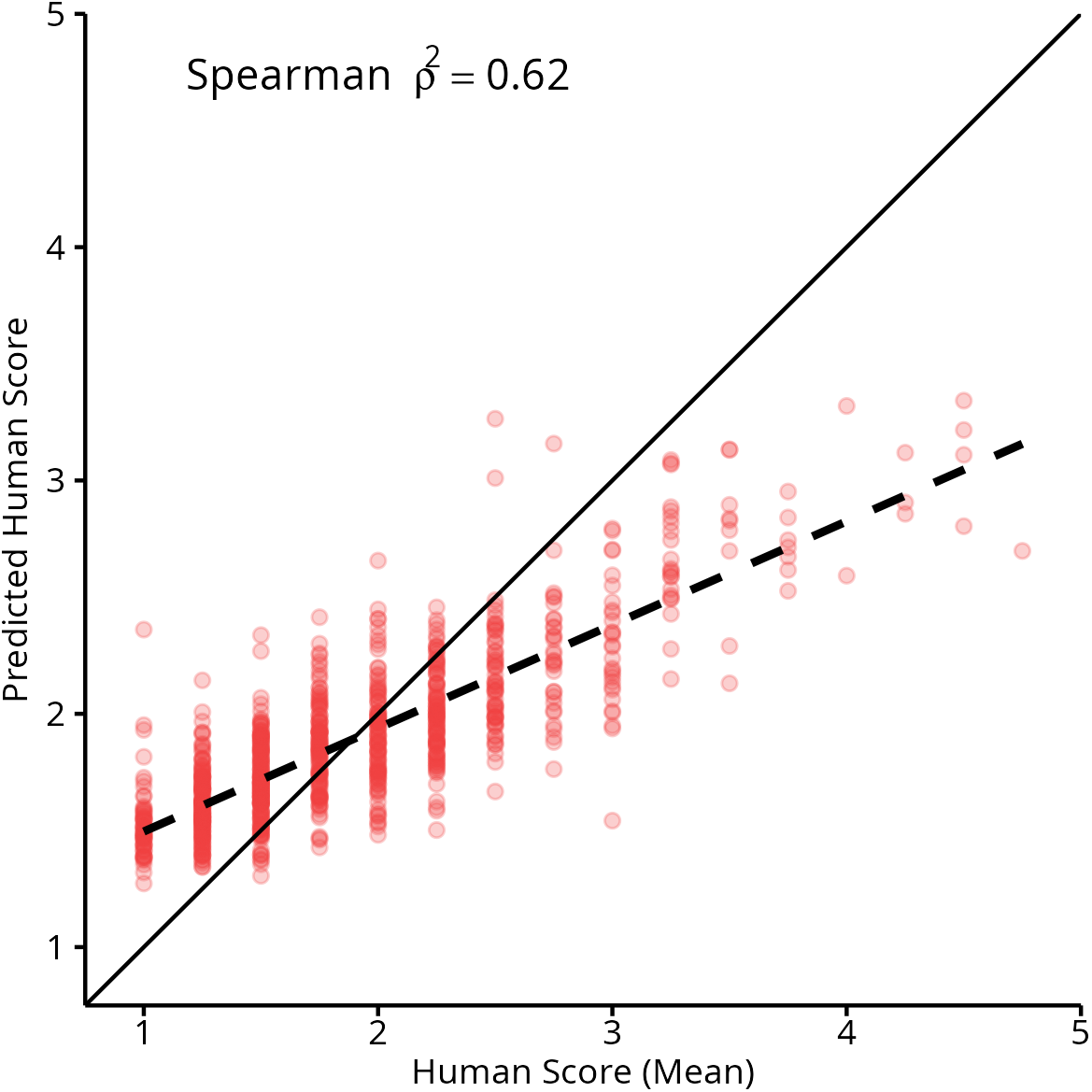
Human disease ratings versus DINOv2 random-forest predictions. Solid line, equality; dashed line, linear fit.

**Fig. S5.**
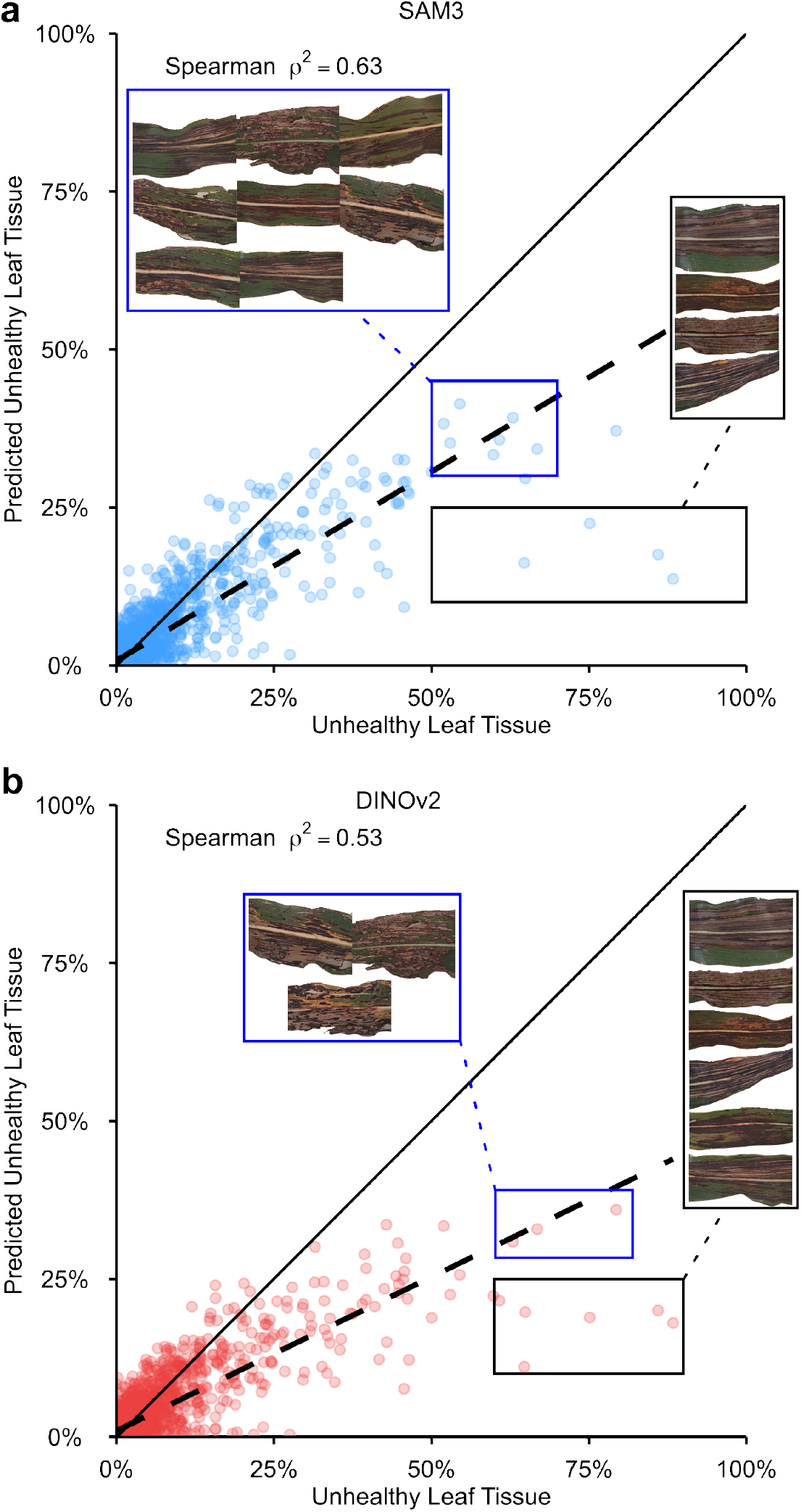
ExG-estimated unhealthy leaf area versus random-forest predictions. **a** SAM3. **b** DINOv2. Solid lines indicate equality; dashed lines show linear fits. Insets correspond to points in matching colored rectangles.

**Fig. S6.**
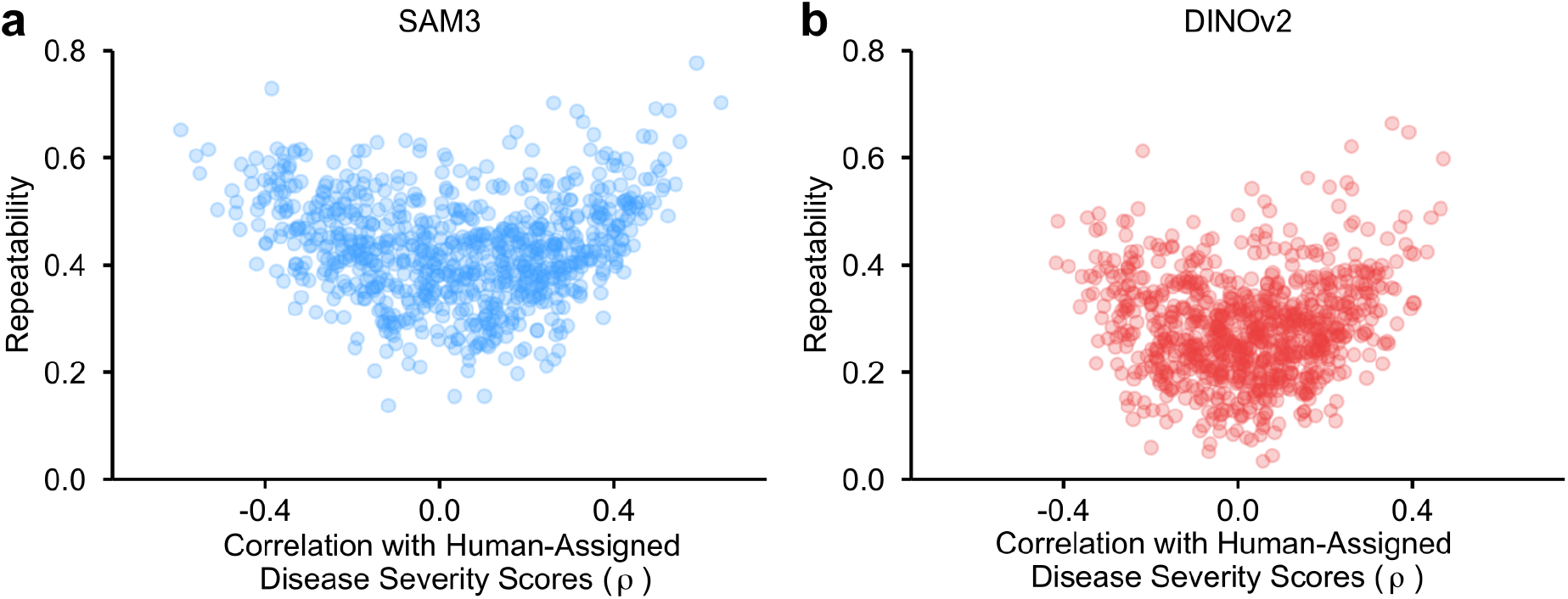
Embedding repeatability versus correlation with human-assigned disease severity scores in Nebraska. Correlations are Spearman *ρ* for SAM3 (**a**) and DINOv2 (**b**).

**Fig. S7.**
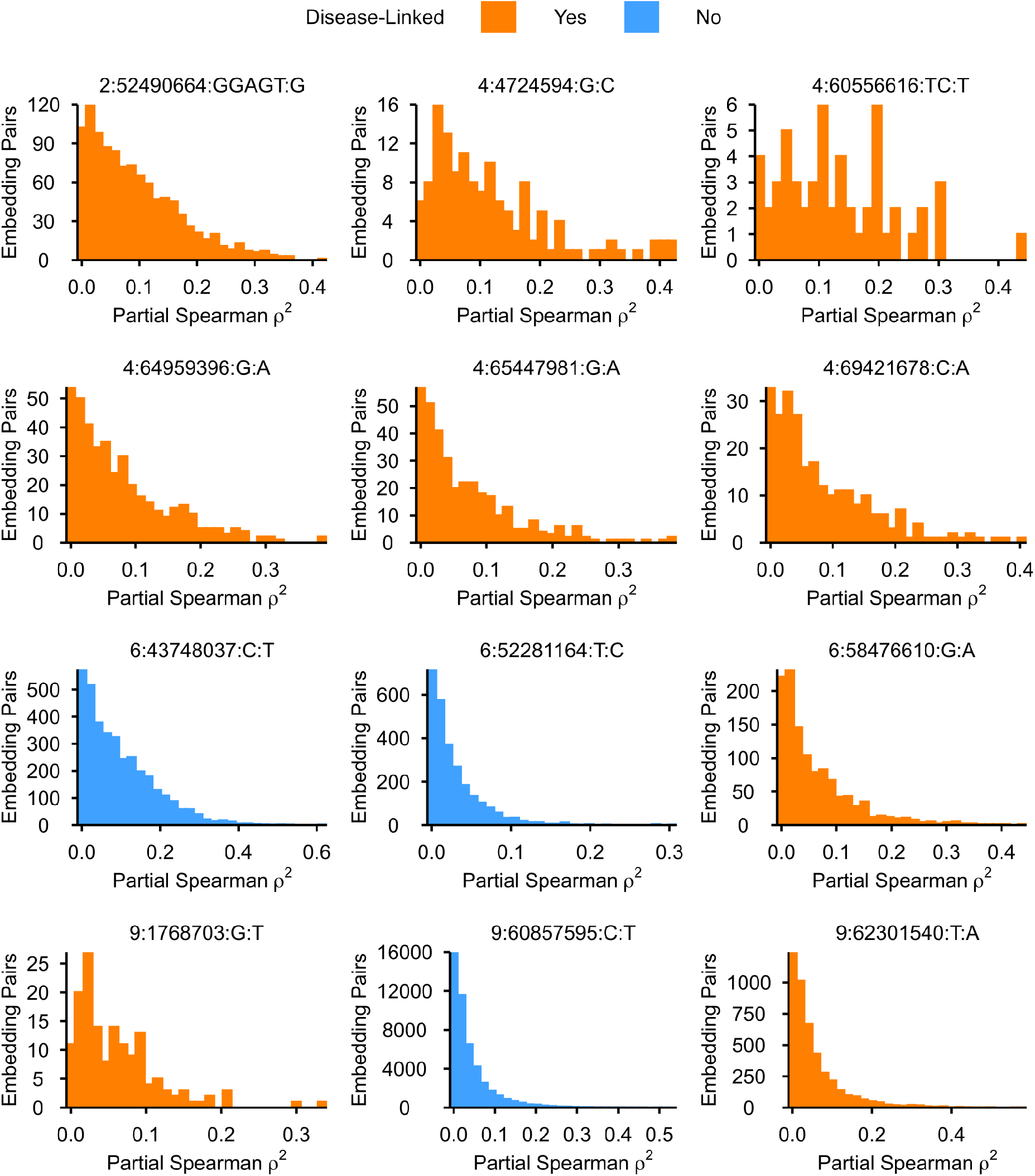
Within-hotspot embedding correlations after covariate adjustment. Squared partial Spearman correlations (*ρ*^2^) for pairs of SAM3 and DINOv2 embeddings associated with each hotspot. Colors distinguish disease-linked hotspots; adjustment is described in Methods.

**Fig. S8.**
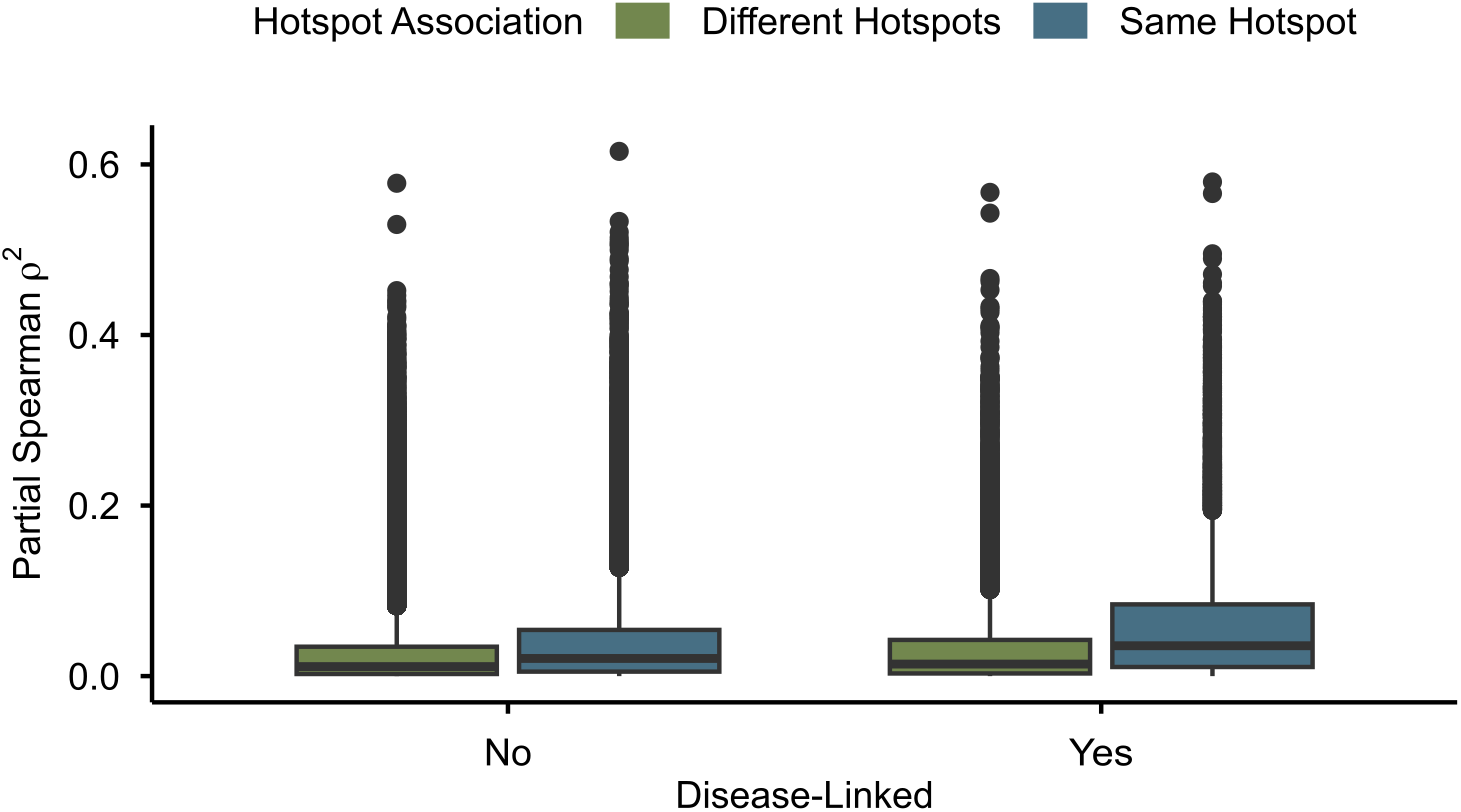
Partial embedding correlations within and between hotspots. Squared partial Spearman correlations (*ρ*^2^) include SAM3 and DINOv2 embedding pairs. Between-hotspot pairs are disease-linked only when both embeddings map to at least one disease-linked hotspot. Pair counts (different/same hotspot): 115,387/50,470 not disease-linked; 26,869/11,082 disease-linked. Adjustment is described in Methods.

**Fig. S9.**
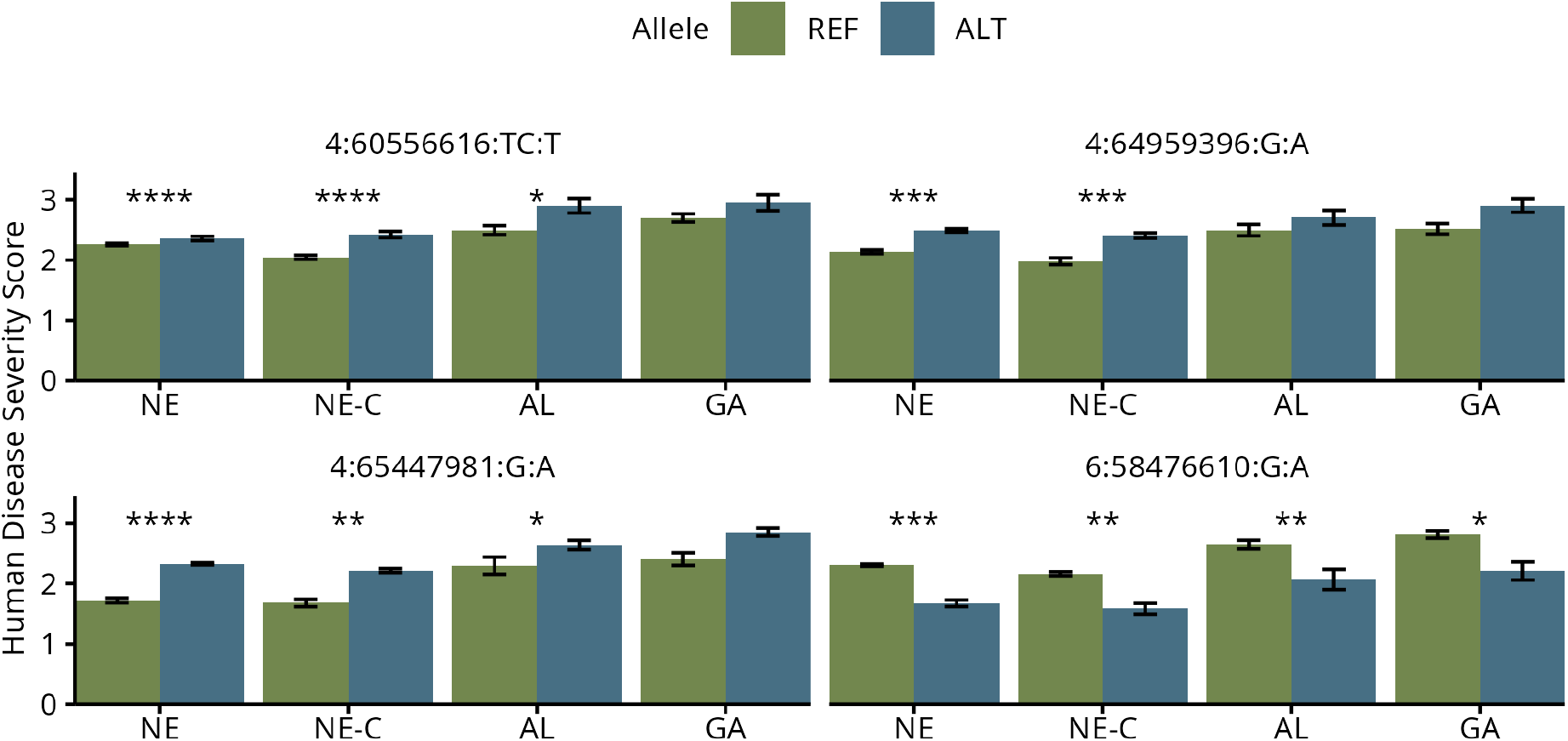
Human disease-score associations across environments. Bars show homozygote BLUE means *±* SE. NE, Nebraska; NE-C, Nebraska genotypes shared across all environments; AL, Alabama; GA, Georgia. Tests include eligible heterozygotes, which are not plotted. Asterisks denote single-marker likelihood-ratio tests: \**p <* 0.05, \*\**p <* 0.01, \*\*\**p <* 0.001, \*\*\*\**p <* 0.0001. Plotted REF/ALT genotype counts, in NE, NE-C, AL, GA order: 4:60556616, 725/144, 188/40, 101/28, 117/26; 4:64959396, 277/245, 77/60, 58/35, 44/43; 4:65447981, 81/786, 38/189, 21/106, 31/111; 6:58476610, 810/40, 213/15, 117/12, 129/15.

**Fig. S10.**
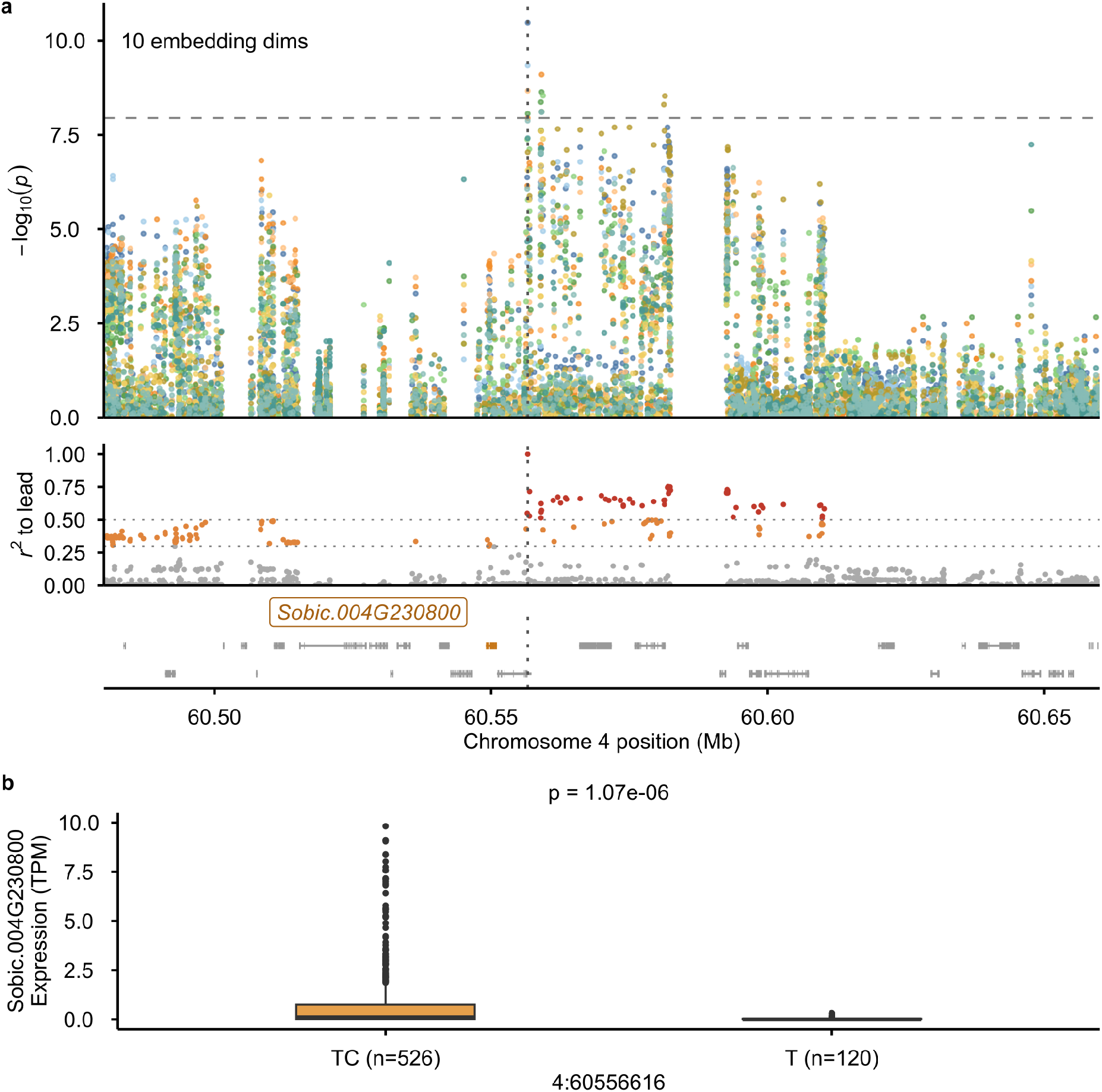
Associations at the UDP-glycosyltransferase candidate *Sobic.004G230800*. **a** SAM3 associations, LD to 4:60556616:TC:T, and gene models. Only embeddings with a genome-wide-significant marker in the displayed region are shown. Horizontal lines mark the GWAS threshold and LD *r*^2^ = 0.3 and 0.5; LD colors are grey (*r*^2^ *≤* 0.3), orange (0.3 *< r*^2^ *≤* 0.5), and red (*r*^2^ *>* 0.5). Vertical lines mark lead markers. Association and LD spans are not fine-mapping confidence intervals. **b** Nebraska 2021 raw-TPM expression, retaining zeros and outliers (*n* = 534 TC/TC and 120 T/T homozygotes). Six heterozygotes and nine TC/TC homozygotes with expression *>* 10 TPM were included in the single-marker likelihood-ratio test but omitted from the plot. Boxes show medians and interquartile ranges; whiskers extend to observations within 1.5*×* IQR.

**Fig. S11.**
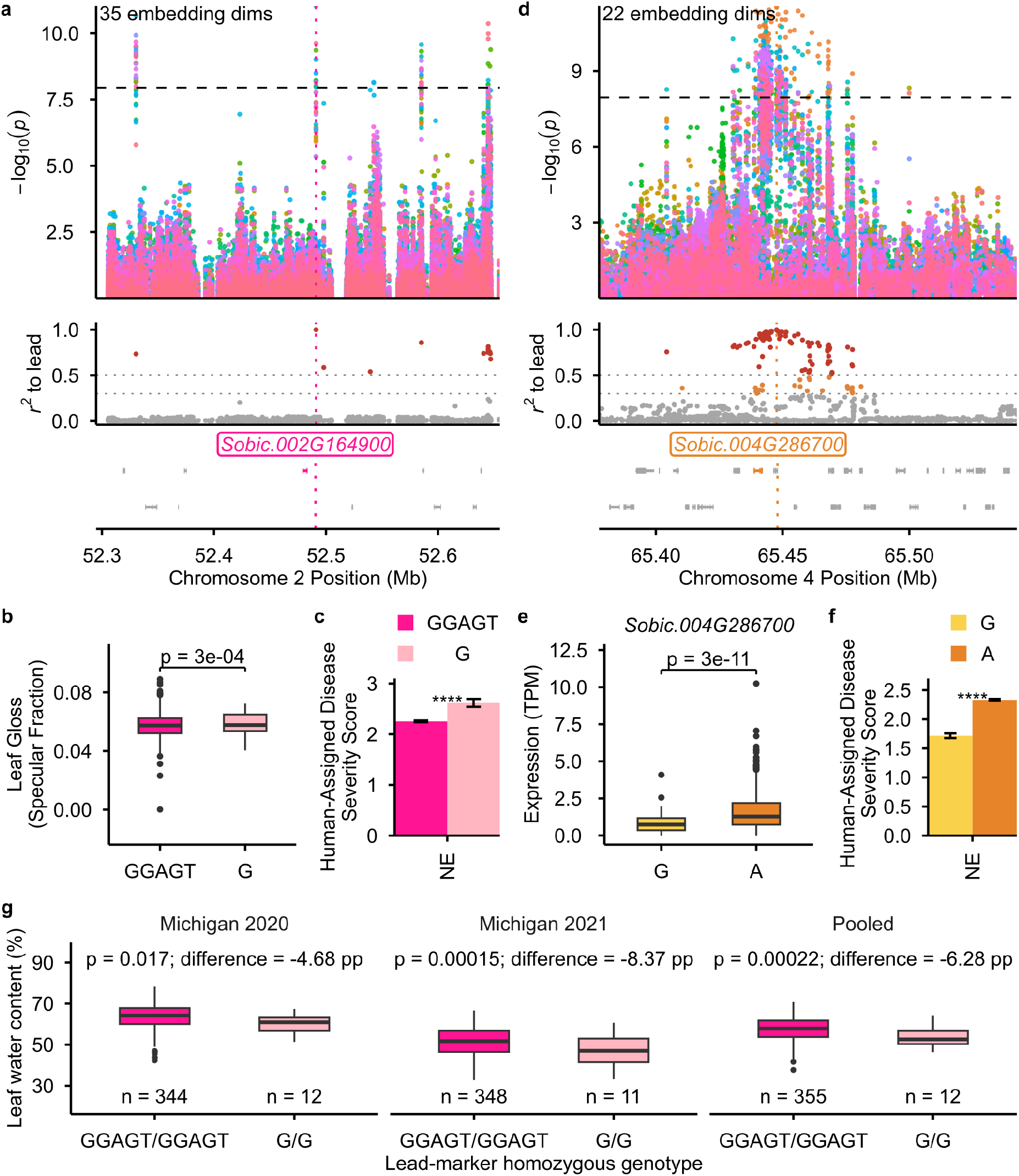
GDSL candidate-gene associations at two disease-linked hotspots. **a, d** SAM3 associations, LD to the lead marker, and gene models; only embeddings with a genome-wide-significant marker in the displayed region are shown. Lines and LD colors follow Fig. S10. **b** Nebraska 2025 leaf specular fraction (*n* = 844 GGAGT and 49 G homozygotes). **c, f** Nebraska 2025 human-assigned disease severity score BLUEs, mean *±* SE (*n* = 841*/*48 GGAGT/G and 80*/*808 G/A homozygotes). **e** Nebraska 2021 raw-TPM expression of *Sobic.004G286700* (*n* = 64 G and 593 A homozygotes), retaining zeros and outliers. **g** Leaf water content [41], 100(fresh *−* dry)*/*fresh; pooled values average each genotype’s environment-specific fractions. Contrasts are alternate-minus-reference homozygote differences in percentage points (pp). Tests retain heterozygotes omitted from plots: two in **c**, three in **e, f**, and one per test in **g**. P-values and asterisks denote single-marker likelihood-ratio tests: \*\*\**p <* 0.001, \*\*\*\**p <* 0.0001. Boxes show medians and interquartile ranges; whiskers extend to observations within 1.5*×* IQR.

**Fig. S12.**
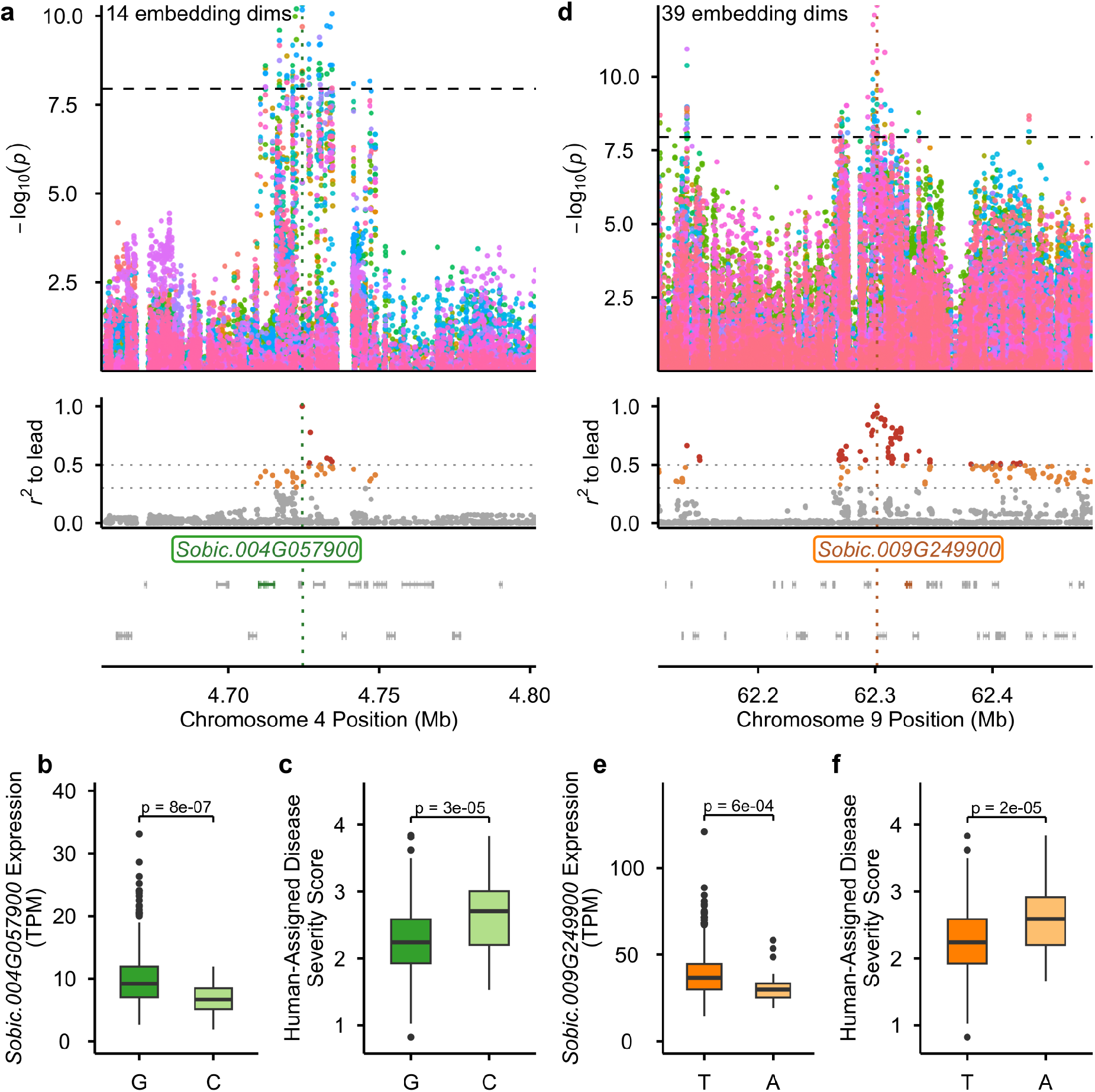
Candidate-gene associations at the chr4:4.7 Mb and chr9:62.2 Mb hotspots. **a, d** SAM3 associations colored by embedding, LD to the lead marker, and gene models. Lines and LD colors follow Fig. S10. **b, e** Nebraska 2021 raw-TPM expression of *Sobic.004G057900* and *Sobic.009G249900*, respectively. **c, f** Nebraska 2025 human-assigned disease severity score BLUEs. Homozygote counts: **b**, 627 G/G and 30 C/C; **c**, 844 G/G and 44 C/C; **e**, 616 T/T and 42 A/A; **f**, 839 T/T and 50 A/A. Tests retain three heterozygotes in **b, c** and two in **e, f**, omitted from plots. Boxes show medians and interquartile ranges; whiskers extend to observations within 1.5*×* IQR. Expression zeros and outliers are retained. Pvalues are from single-marker likelihood-ratio tests.

**Fig. S13.**
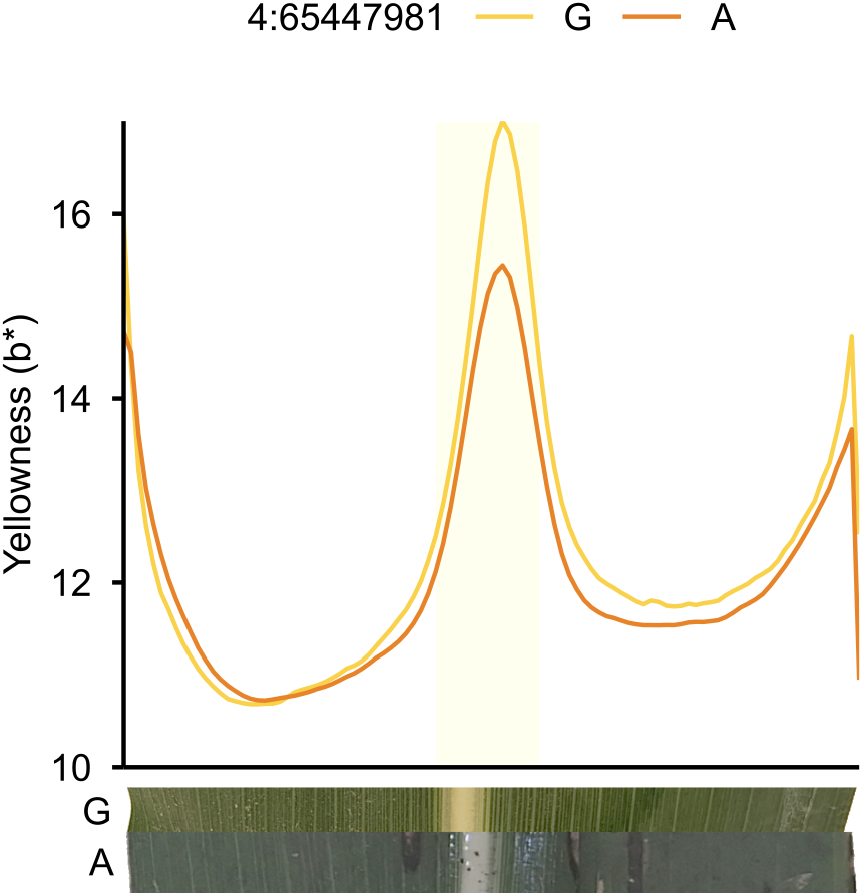
Leaf yellowness profiles by 4:65447981 allele in Nebraska 2025. Lines show genotype-averaged CIELAB b* across 100 leaf-width bins; shading marks the 14 central bins containing the midrib. Insets show representative leaf slices. *n* = 81 G and 786 A genotypes.

**Fig. S14.**
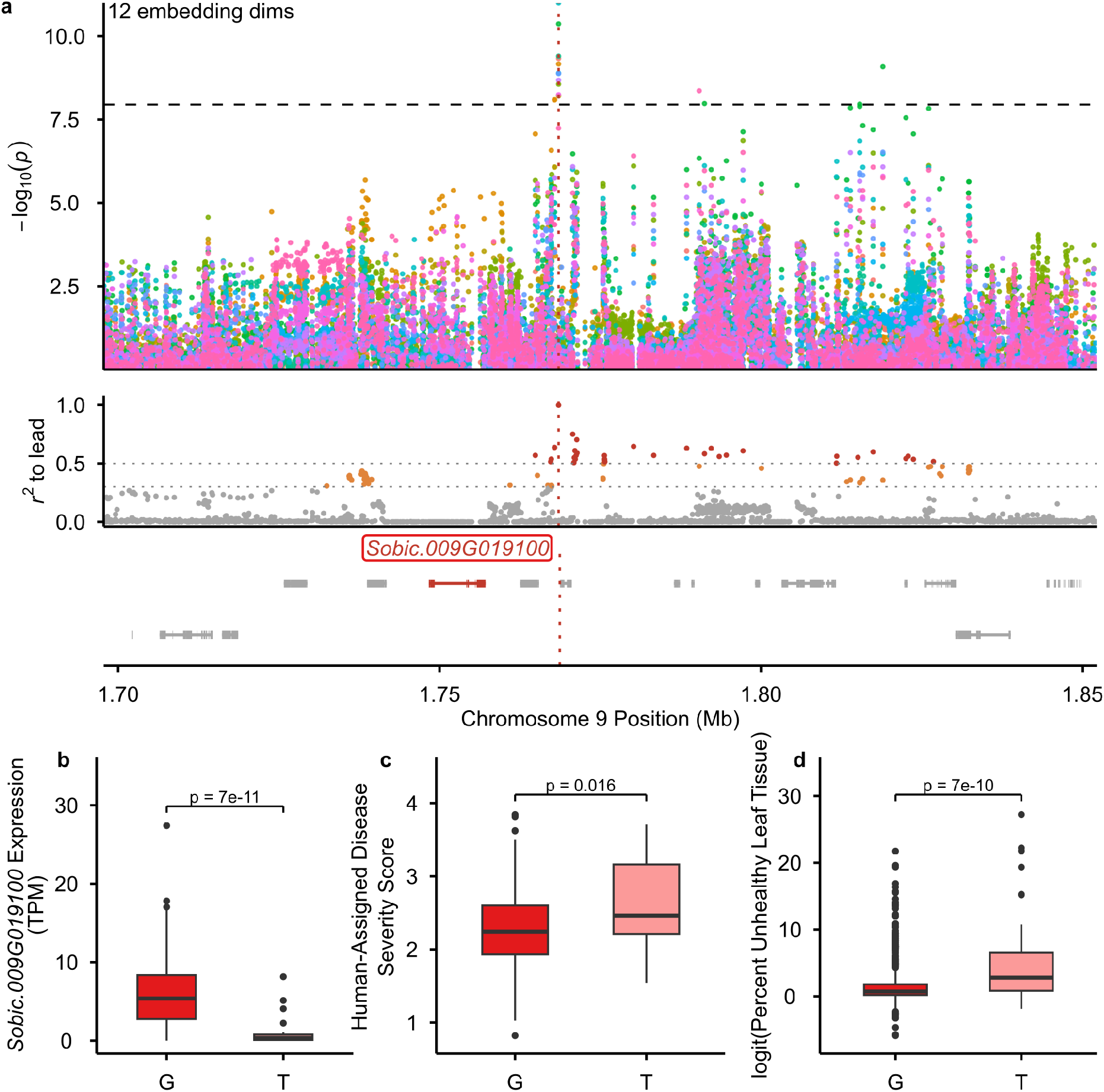
Associations at the LysM candidate gene *Sobic.009G019100*. **a** SAM3 associations, LD to 9:1768703:G:T, and gene models. Lines and LD colors follow Fig. S10. **b** Nebraska 2021 raw-TPM expression (*n* = 624 G/G and 36 T/T), retaining zeros and outliers. **c, d** Nebraska 2025 human-assigned disease severity score and logit(unhealthy leaf area) BLUEs, respectively (*n* = 852 G/G and 39 T/T each). No heterozygotes were observed. Boxes show medians and interquartile ranges; whiskers extend to observations within 1.5*×* IQR. P-values are from single-marker likelihood-ratio tests.

**Fig. S15.**
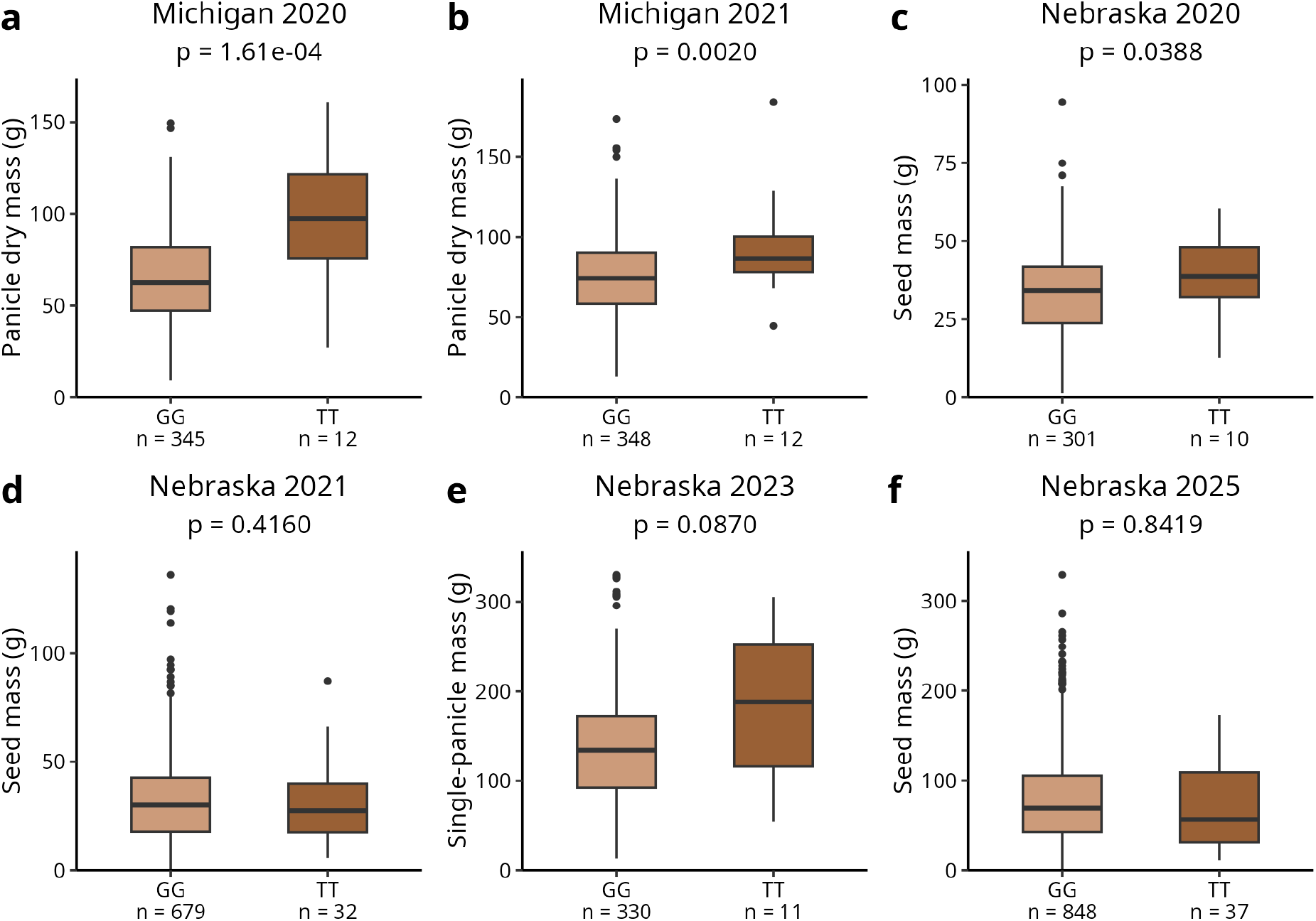
Grain- and panicle-mass associations with Chr09:1768703:G:T across six environments. Plots show genotype means; heterozygotes were excluded. Boxes show medians and interquartile ranges; whiskers extend to observations within 1.5*×* IQR. Nominal p-values are from single-marker likelihood-ratio tests.

**Fig. S16.**
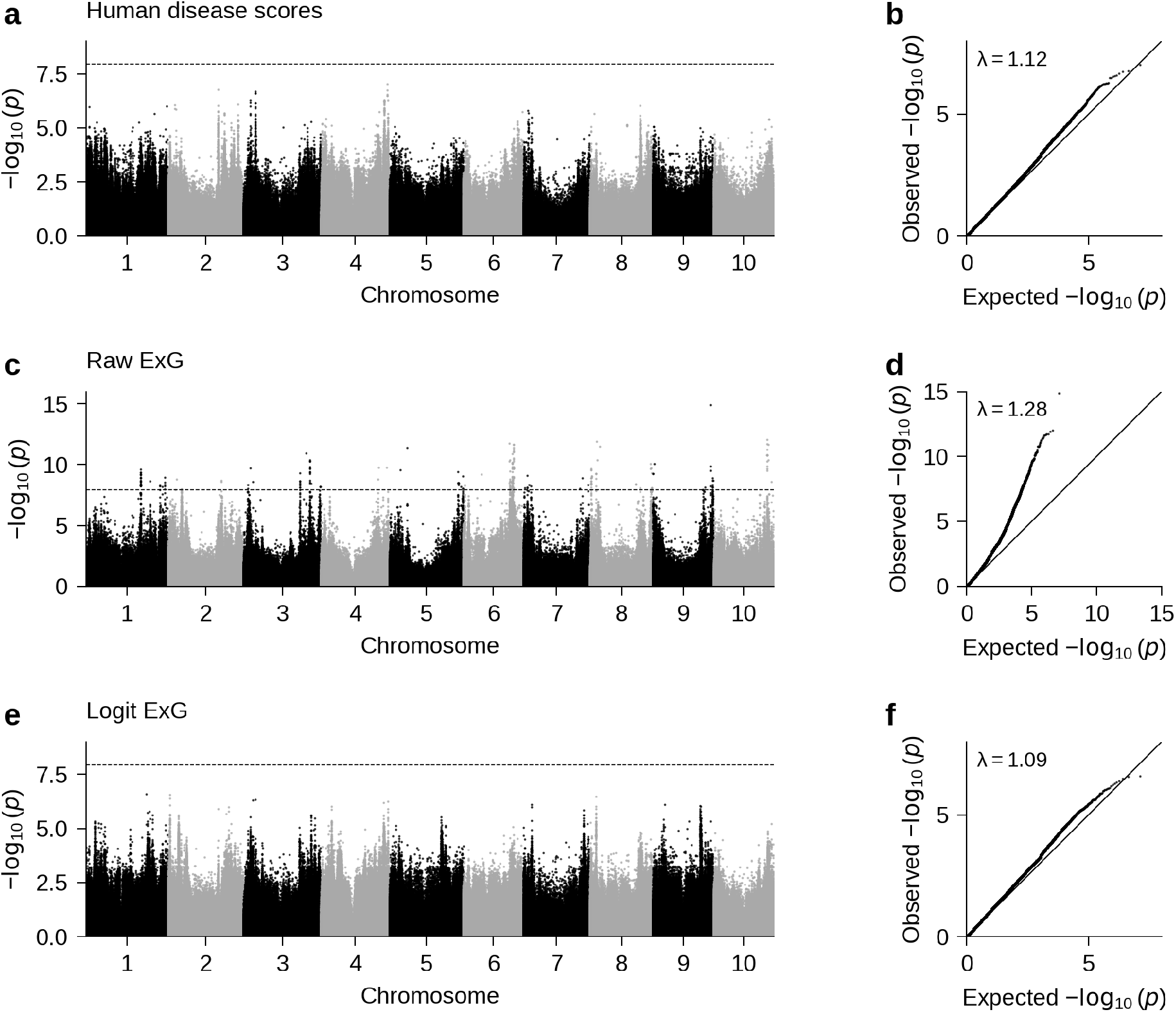
Genome-wide associations for human-assigned and ExG-based disease scores. **a, b** Human-assigned disease severity scores. **c, d** Untransformed ExG estimates of unhealthy leaf area. **e, f** Logit-transformed ExG estimates of unhealthy leaf area. Manhattan plots are shown at left and quantile–quantile plots at right. Each GWAS tested 6,422,975 markers in 891 Nebraska 2025 genotypes using the BLUE and PANICLE mixed-model procedures described in Methods, with leave-one-chromosome-out kinship and with five genetic principal components, leaf area, and flowering time as covariates. Dashed horizontal lines mark the effective-Bonferroni threshold (*p* = 0.05*/*4,446,367; *−* log10 *p ≈* 7.95). Diagonal lines in the quantile–quantile plots indicate the null expectation; *λ* denotes the genomic inflation factor calculated from the median one-degree-of-freedom chi-square statistic.

**Fig. S17.**
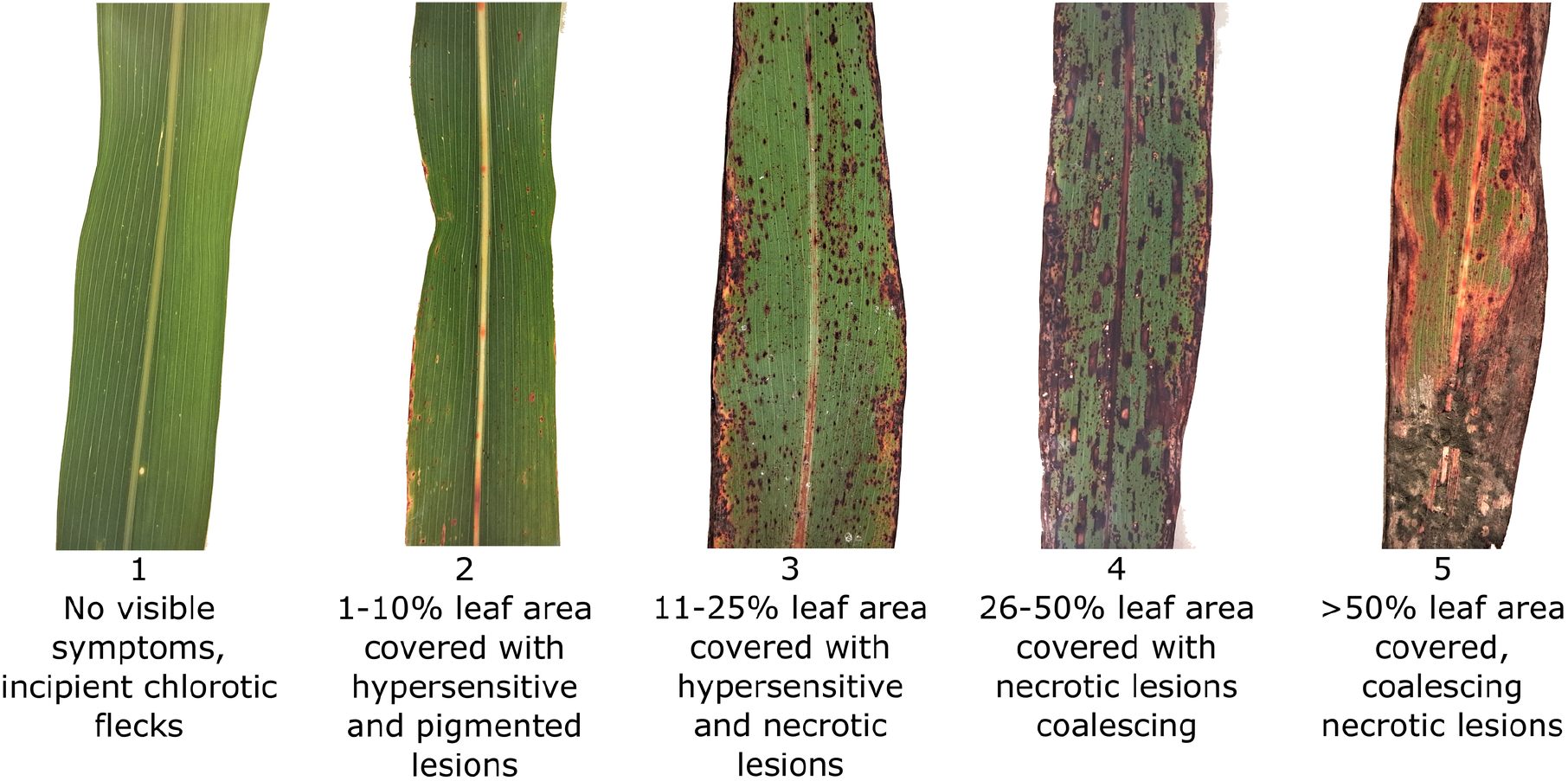
Scale used for human-assigned anthracnose severity scores. Adapted from Thakur [14].

**Table S1.**
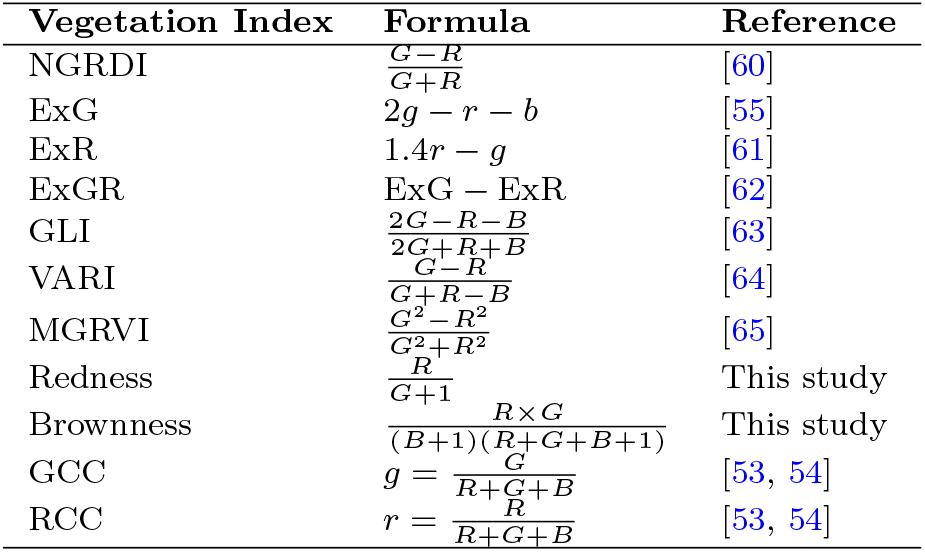
RGB-based vegetation indices evaluated in this study.

**Table S2.** Independently defined DINOv2 embedding GWAS hotspots. Peak bounds are rounded to 0.1 Mb; marker positions are in base pairs (BTx623 v5). Counts are distinct associated DINOv2 embeddings per merged hotspot, not per window. Peak markers have the strongest embedding association within each hotspot.

| Peak | Peak marker | Embeddings |
| --- | --- | --- |
| Chr06:51.2–51.3 | Chr06:51301175 | 11 |
| Chr06:51.4–51.5 | Chr06:51436451 | 10 |
| Chr06:51.7–51.9 | Chr06:51804054 | 14 |
| Chr06:51.9–52.6 | Chr06:52190357 | 64 |
| Chr06:58.5–58.7 | Chr06:58572334 | 18 |
| Chr09:59.8–61.8 | Chr09:60868989 | 173 |
| Chr09:61.8–62.2 | Chr09:61927561 | 39 |
| Chr09:62.2–62.4 | Chr09:62272432 | 11 |

**Table S3.** Cross-environment replication of SAM3 embedding associations. Embeddings mapping to multiple hotspots are counted separately. NE-common eligible pairs are testable and significant (*p <* 0.05) in Nebraska genotypes shared across all three environments. Replication requires nominal *p <* 0.05 in Alabama (AL), Georgia (GA), either (*≥* 1), or both. Eligibility percentages use total pairs; replication percentages use eligible pairs. Dashes denote undefined percentages when no pairs are eligible.

| Hotspot<br>(Chr.:Mb) | Total<br>pairs | NE-common<br>eligible, n (%)<br>total) | AL, n (%)<br>eligible) | GA, n (%)<br>eligible) | $\geq 1$, n (%)<br>eligible) | Both, n<br>(%<br>eligible) |
| --- | --- | --- | --- | --- | --- | --- |
| 2:52.3 | 40 | 13 (32.5) | 0 (0.0) | 4 (30.8) | 4 (30.8) | 0 (0.0) |
| 4:4.7 | 14 | 2 (14.3) | 0 (0.0) | 0 (0.0) | 0 (0.0) | 0 (0.0) |
| 4:60.5 | 10 | 10 (100.0) | 0 (0.0) | 2 (20.0) | 2 (20.0) | 0 (0.0) |
| 4:64.9 | 24 | 21 (87.5) | 5 (23.8) | 7 (33.3) | 10 (47.6) | 2 (9.5) |
| 4:65.4 | 22 | 15 (68.2) | 2 (13.3) | 4 (26.7) | 5 (33.3) | 1 (6.7) |
| 4:69.4 | 19 | 2 (10.5) | 0 (0.0) | 0 (0.0) | 0 (0.0) | 0 (0.0) |
| 6:44.1 | 75 | 74 (98.7) | 4 (5.4) | 23 (31.1) | 23 (31.1) | 4 (5.4) |
| 6:52.1 | 20 | 17 (85.0) | 1 (5.9) | 6 (35.3) | 7 (41.2) | 0 (0.0) |
| 6:58.5 | 30 | 25 (83.3) | 10 (40.0) | 17 (68.0) | 17 (68.0) | 10 (40.0) |
| 9:1.7 | 12 | 0 (0.0) | 0 (—) | 0 (—) | 0 (—) | 0 (—) |
| 9:60.8 | 140 | 106 (75.7) | 7 (6.6) | 25 (23.6) | 29 (27.4) | 3 (2.8) |
| 9:62.2 | 54 | 22 (40.7) | 0 (0.0) | 2 (9.1) | 2 (9.1) | 0 (0.0) |
| **All hotspots** | **460** | **307 (66.7)** | **29 (9.4)** | **90 (29.3)** | **99 (32.2)** | **20 (6.5)** |

**Table S4.** PheWAS sample sizes. Entries count analyzed genotypes, not individual plants; dashes indicate combinations not analyzed. NE, Nebraska; MI, Michigan; AL, Alabama.

| **Trait** | **NE**<br>**2020** | **MI**<br>**2020** | **NE**<br>**2021** | **MI**<br>**2021** | **AL**<br>**2022** | **NE**<br>**2023** | **NE**<br>**2025** |
| --- | --- | --- | --- | --- | --- | --- | --- |
| Days To Flower | 324 | 365 | 836 | 836 | 366 | — | 899 |
| Plant Height | — | 365 | — | 369 | — | — | 902 |
| Flag Leaf Length | 322 | — | — | — | — | — | — |
| Flag Leaf Width | 322 | — | — | — | — | — | — |
| Leaf Length | 323 | 365 | 479 | 369 | — | — | — |
| Leaf Width | 323 | 365 | 479 | 369 | — | — | — |
| Leaf Number | 323 | — | 453 | — | — | — | 904 |
| Median Leaf Angle | 324 | — | — | — | — | — | — |
| Tillers Present | 322 | 365 | 485 | 369 | — | — | — |
| Stem Diameter Lower | 322 | 365 | 361 | 369 | — | — | — |
| Stem Diameter Upper | 322 | — | 361 | — | — | — | — |
| Panicles Per Plant | — | — | 766 | — | — | — | — |
| Rachis Length | 322 | — | 315 | — | — | — | — |
| Rachis Diameter Lower | 322 | — | 315 | — | — | — | — |
| Rachis Diameter Upper | 322 | — | 315 | — | — | — | — |
| Primary Branch Count | 322 | — | 315 | — | — | — | — |
| Branch Internode Length | 322 | — | 315 | — | — | — | — |
| Seed Mass | 318 | — | 725 | — | — | — | 899 |
| Seed Protein | 314 | — | 791 | — | 365 | 309 | — |
| Seed Oil | 314 | — | 782 | — | 361 | 309 | — |
| Seed Ash | 314 | — | 791 | — | 365 | 309 | — |
| Seed Starch | 314 | — | 791 | — | 365 | 309 | — |
| Chlorophyll Concentration | 314 | — | — | — | — | 351 | 903 |
| Flag Leaf Height | 323 | 365 | 837 | 369 | 366 | — | 901 |
| Single Panicle Mass | — | — | — | — | — | 349 | — |
| Seed Area | — | — | 806 | — | — | — | 310 |
| Seed Length | — | — | 806 | — | — | — | 310 |
| Seed Width | — | — | 806 | — | — | — | 310 |
| Seed Blue Intensity | — | — | 805 | — | — | — | 310 |
| Seed Green Intensity | — | — | 805 | — | — | — | 310 |
| Seed Red Intensity | — | — | 805 | — | — | — | 310 |
| Panicle Length | — | 365 | — | 369 | — | — | 901 |
| Final Green Leaves | — | 365 | — | 369 | — | — | — |
| Stem Volume | 322 | 365 | 360 | 369 | — | — | — |
| Single Plant Fresh Weight | — | 365 | — | 369 | — | — | — |
| Single Plant Dry Weight | — | 365 | — | 368 | — | — | — |
| Single Plant Stem Fresh Weight | — | 365 | — | 368 | — | — | — |
| Single Plant Stem Dry Weight | — | 365 | — | 368 | — | — | — |
| Single Plant Leaf Fresh Weight | — | 365 | — | 368 | — | — | — |
| Single Plant Leaf Dry Weight | — | 365 | — | 368 | — | — | — |
| Single Plant Panicle Fresh Weight | — | 365 | — | 368 | — | — | — |
| Single Plant Panicle Dry Weight | — | 365 | — | 368 | — | — | — |
| Total Plot Fresh Weight | — | 363 | — | 369 | — | — | — |
| Total Plot Dry Weight | — | 363 | — | 369 | — | — | — |
| Thousand Kernel Weight | — | — | — | — | — | — | 304 |
| Leaf Thickness | 323 | 365 | — | — | — | — | 904 |
| Leaf Area Index | — | 365 | — | — | — | — | — |
| Specific Leaf Area | 314 | 365 | — | — | — | — | — |

**Information S1** STL files and parts list for portable leaf imaging chamber

**Information S2** Illustrated image collection protocol.

**Information S3** Source code for custom Python web app used for human scoring of disease severity in images.

## Notes

https://doi.org/10.5061/dryad.cfxpnvxpb

https://github.com/jschnable/SorghumLeafEmbeddings/

